# Adaptation of the *Mycobacterium tuberculosis* transcriptome to biofilm growth

**DOI:** 10.1101/2023.07.18.549484

**Authors:** Madison A. Youngblom, Tracy M. Smith, Caitlin S. Pepperell

## Abstract

*Mycobacterium tuberculosis* (*M. tb*), the causative agent of tuberculosis (TB), is a leading global cause of death from infectious disease. Biofilms are increasingly recognized as a relevant growth form during *M. tb* infection and may impede treatment by enabling bacterial drug and immune tolerance. *M. tb* has a complicated regulatory network that has been well-characterized for many relevant disease states, including dormancy and hypoxia. However, despite its importance, our knowledge of the genes and pathways involved in biofilm formation is limited. Here we characterize the biofilm transcriptomes of fully virulent clinical isolates and find that the regulatory systems underlying biofilm growth vary widely between strains and are also distinct from regulatory programs associated with other environmental cues. We used experimental evolution to investigate changes to the transcriptome during adaptation to biofilm growth and found that the application of a uniform selection pressure resulted in loss of strain-to-strain variation in gene expression, resulting in a more uniform biofilm transcriptome. The adaptive trajectories of transcriptomes were shaped by the genetic background of the *M. tb* population leading to convergence on a sub-lineage specific transcriptome. We identified widespread upregulation of non-coding RNA (ncRNA) as a common feature of the biofilm transcriptome and hypothesize that ncRNA function in genome-wide modulation of gene expression, thereby facilitating rapid regulatory responses to new environments. These results reveal a new facet of the *M. tb* regulatory system and provide valuable insight into how *M. tb* adapts to new environments.

**Importance:** Understanding mechanisms of resistance and tolerance in *Mycobacterium tuberculosis* (*M. tb*) can help us develop new treatments that capitalize on *M. tb*’s vulnerabilities. Here we used transcriptomics to study both the regulation of biofilm formation in clinical isolates as well as how those regulatory systems adapt to new environments. We find that closely related clinical populations have diverse strategies for growth under biofilm conditions, and that genetic background plays a large role in determining the trajectory of evolution. These results have implications for future treatment strategies that may be informed by our knowledge of the evolutionary constraints on strain(s) from an individual infection. This work provides new information about the mechanisms of biofilm formation in *M. tb* and outlines a framework for population level approaches for studying bacterial adaptation.

## Introduction

*Mycobacterium tuberculosis* (*M. tb*), the causative agent of tuberculosis (TB), is a globally distributed and highly prevalent pathogen. *M. tb* is uniquely adapted to its environment within human tissues and is adept at evading both antibiotic treatment and the immune system, resulting in recalcitrant, difficult to treat infections. Increasing drug resistance among populations of *M. tb* is a global health concern: in 2021 there were over 10 million new infections, ∼500,000 of which were multi-drug resistant (*1*). Our ability to treat infections in the face of ever-worsening antibiotic resistance hinges on our understanding of how *M. tb* responds and adapts to the various selective pressures encountered during infection and transmission.

*M. tb*’s regulatory system is highly interconnected (*2*, *3*) and is structured such that genome-wide patterns of gene expression can be changed rapidly via master regulatory elements (*2*, *4*). Features of this regulatory system have been well-studied under a number of conditions relevant to *M. tb* pathogenesis including dormancy (*5*) and hypoxia (*6*–*8*). These studies have outlined regulatory programs used by *M. tb* to adapt to these conditions including the DosR regulon, which is responsible for coordinating gene expression during non-replicative dormancy. Non-replicative dormancy is thought to allow *M. tb* to persist during latent infection.

Studies of human autopsies (*9*–*11*) and animal models (*12*) have identified aggregated *M. tb* cells that resemble biofilms, suggesting that biofilms are an important growth form for *M. tb* during natural infection. Biofilms contribute to infection persistence via immune evasion and drug tolerance (*12*–*15*). Yet, we know very little about biofilm formation, particularly the regulatory processes governing this growth form. A single transcriptomics study of *M. tb* pellicle biofilms has been published, in which the authors identified a role for isonitrile lipopeptide (INLP) in biofilm growth (*16*). In a recent paper we describe adaptation of *M. tb* clinical isolates to biofilm growth, using an evolve and re-sequence approach. We discovered that the mutational pathway to enhanced biofilm growth varied between clinical isolates (*17*). These results led us to hypothesize that the regulatory mechanisms underlying pellicle biofilm formation exhibit similar variation.

Here we use experimental evolution and high-coverage transcriptomics to delineate transcriptional responses to pellicle biofilm growth across six clinical strains belonging to two *M. tb* sub-lineages of lineage 4 (L4.9 and L4.2). We further quantify changes to the transcriptome after application of a uniform selective pressure for pellicle growth. We found these relatively closely related clinical strains of *M. tb* to have diverse transcriptional responses to biofilm growth. The application of a uniform selective pressure for pellicle growth caused the strains to converge on a shared transcriptome characterized by wide scale downregulation of gene expression. Within this global pattern we could differentiate two subtypes that were associated with bacterial sub-lineage. Strains from the same sub-lineage evolved a characteristic transcriptome, although the mutational path to this shared transcriptome varied from strain to strain. This implies that pre-existing genetic differences, even among closely related strains, have a strong impact on transcriptome adaptation. We further found that the *M. tb* biofilm transcriptome is characterized by widespread upregulation of non-coding RNA (ncRNA) and hypothesize that expression of ncRNA contributes to genome-wide gene expression modulation.

## Results

### RNA sequencing of clinical and experimental M. tb populations

We previously passaged six clinical populations of *M. tb* (isolated from sputum and not subjected to single-colony passage) under selective pressure to grow as pellicle biofilms as described in T. M. Smith et al., 2022. We identified phenotypic changes during adaptation to biofilm growth including changes in cell morphology, increased extracellular matrix (ECM) production and alterations of growth rate. We also identified genetic changes accompanying enhanced biofilm growth that varied according to the genetic background of the population (*17*).

Here, we sought to assess how serial passage affected the transcriptome. In order to investigate the impacts of biofilm adaptation on global gene expression, we performed RNA sequencing on the ancestral and evolved populations grown under biofilm and planktonic conditions (Figure 1). Two biological replicates were sequenced for each combination of population, genotype, and condition – excepting ancestral planktonic MT49, for which only one replicate was sequenced (Table S1). We extracted between 0.4 and 7 μg of total RNA for each sample and sequenced to high coverage: between 50-120 million reads per sample (Table S1). Sequencing reads were quality checked, trimmed, and aligned to the *M. tb* H37Rv reference genome (NCBI accession NC_000962.3). For calculating expression counts we used HTSeq (*18*) and a custom annotation file (see Methods). Differential expression analysis between conditions (biofilm and planktonic) and across passaging (ancestral and evolved) was performed in R using DESeq2 (*19*).

**Figure 1:**
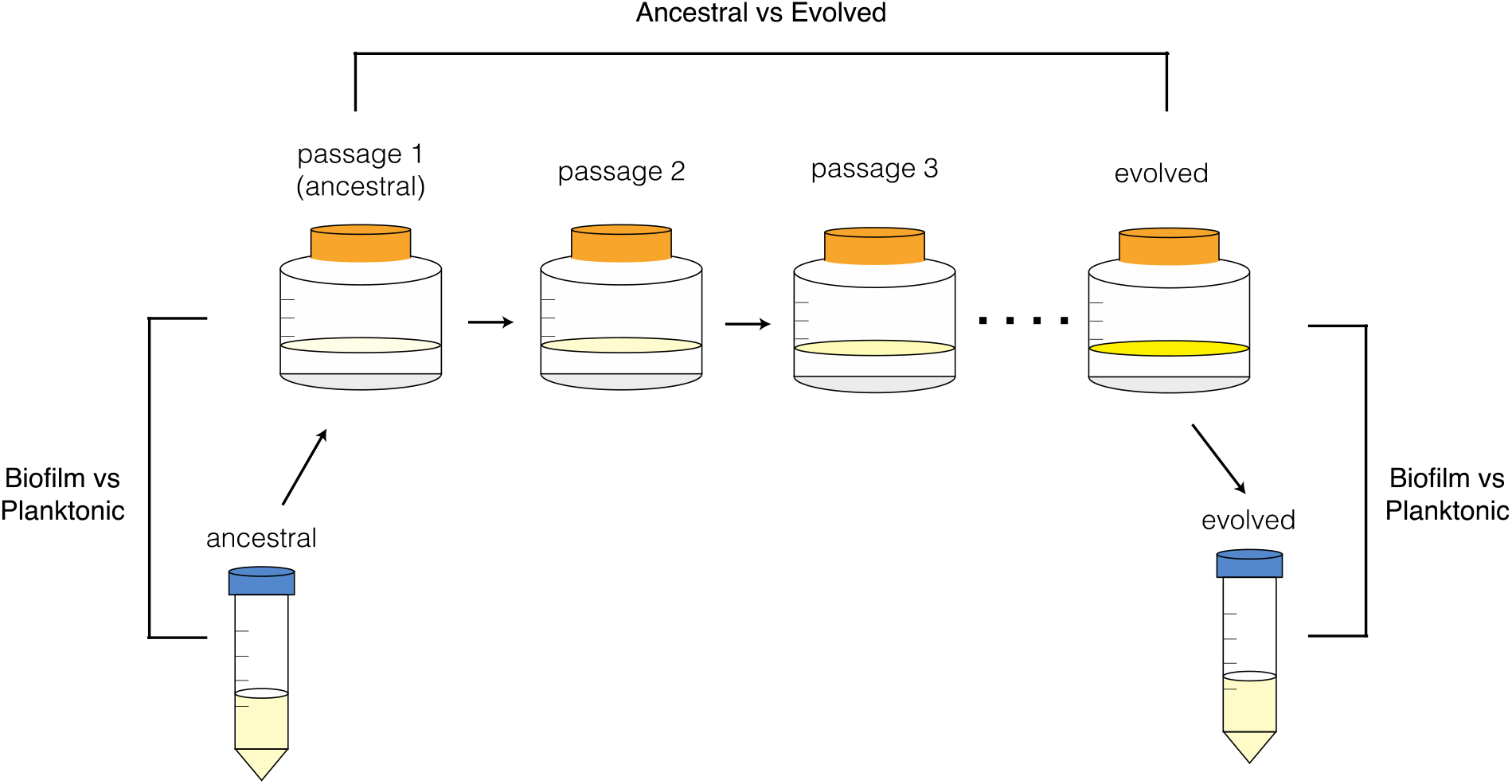
Protocol for the serial passage of *M. tb* pellicle biofilms and subsequent transcriptomic comparisons. Ancestral (clinical) populations were grown in planktonic cultures, then grown as pellicle biofilms (passage 1) before passaging 8-12 times. This passaged (evolved) population was taken from a biofilm culture and grown planktonically again. We performed RNA sequencing on all ancestral and evolved populations, grown under both biofilm and planktonic conditions.

### Genetic background shapes the biofilm transcriptome

We compared the transcriptomes of ancestral bacterial populations grown under biofilm and planktonic conditions. Genetic background appears to have a substantial impact on these transcriptomes, which were consistent among biological replicates but differentiated between populations (Figure 2), with intermingling of planktonic and biofilm samples (Figure 3B, S1A). Differences among genetic backgrounds were also evident in the transcriptomic response to biofilm growth. When analyzed individually, our ancestral populations had between ∼ 500-3000 DEGs in comparisons of biofilm with planktonic conditions (Figure 2, Supplementary Data 1). However, we detected only 143 DEGs with log2 fold changes > +/-2 when all populations were analyzed together (Figure 3C), compared to an average of 625 genes per population with log2 fold changes > +/-2 when populations were analyzed separately (Supplementary Data 1). There is very little overlap in DEGs between populations: only 4% of downregulated genes are shared by at least 5 populations and no upregulated genes are shared across more than 4 populations (Figure 3D). To assess whether the effect of genetic background was reflective of phenotypic differences between ancestral populations, we overlayed wet weight measurements (the simplest marker of biofilm phenotype which measures the total biomass of the biofilm) onto normalized, log-transformed gene expression (Figure S1B). We did not identify a simple relationship between transcriptomic signature and biofilm mass, as samples with similar biomass were spread across transcriptomic space summarized in the PCA. Overall, our results indicate the biofilm transcriptome is affected by subtle differentiation between genetic backgrounds separated by just 10s-100s of SNPs (Figure 2).

**Figure 2:**
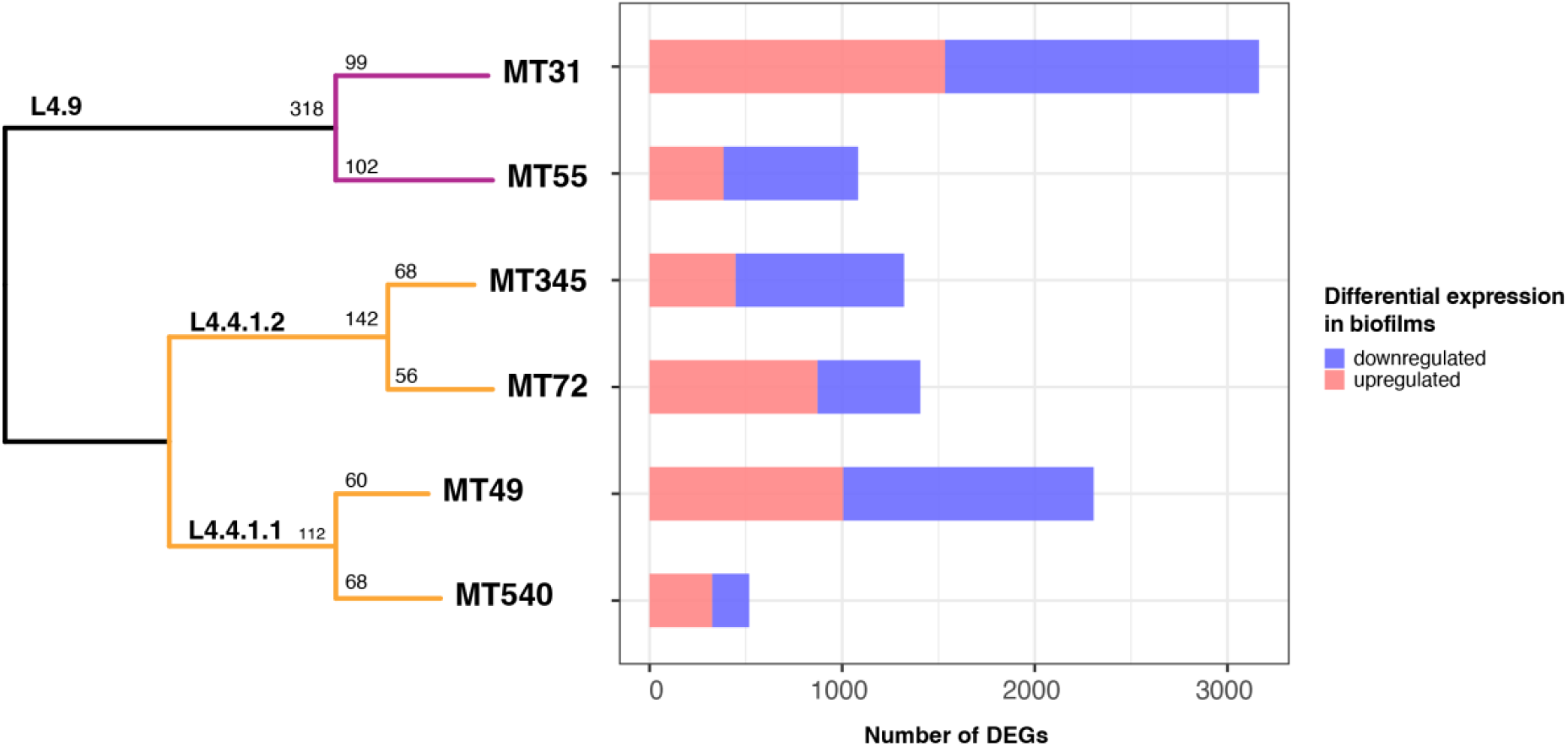
Differentiation of transcriptomes between closely related populations. Left: Whole-genome sequence phylogeny of ancestral populations used for passaging. Populations fall into two major L4 sub-lineages: L4.9 (purple) and L4.4 (orange) with SNP distances shown along each branch. Right: Number of significantly differentially expressed genes (DEGs) between biofilm and planktonic conditions for each ancestral population.

**Figure 3:**
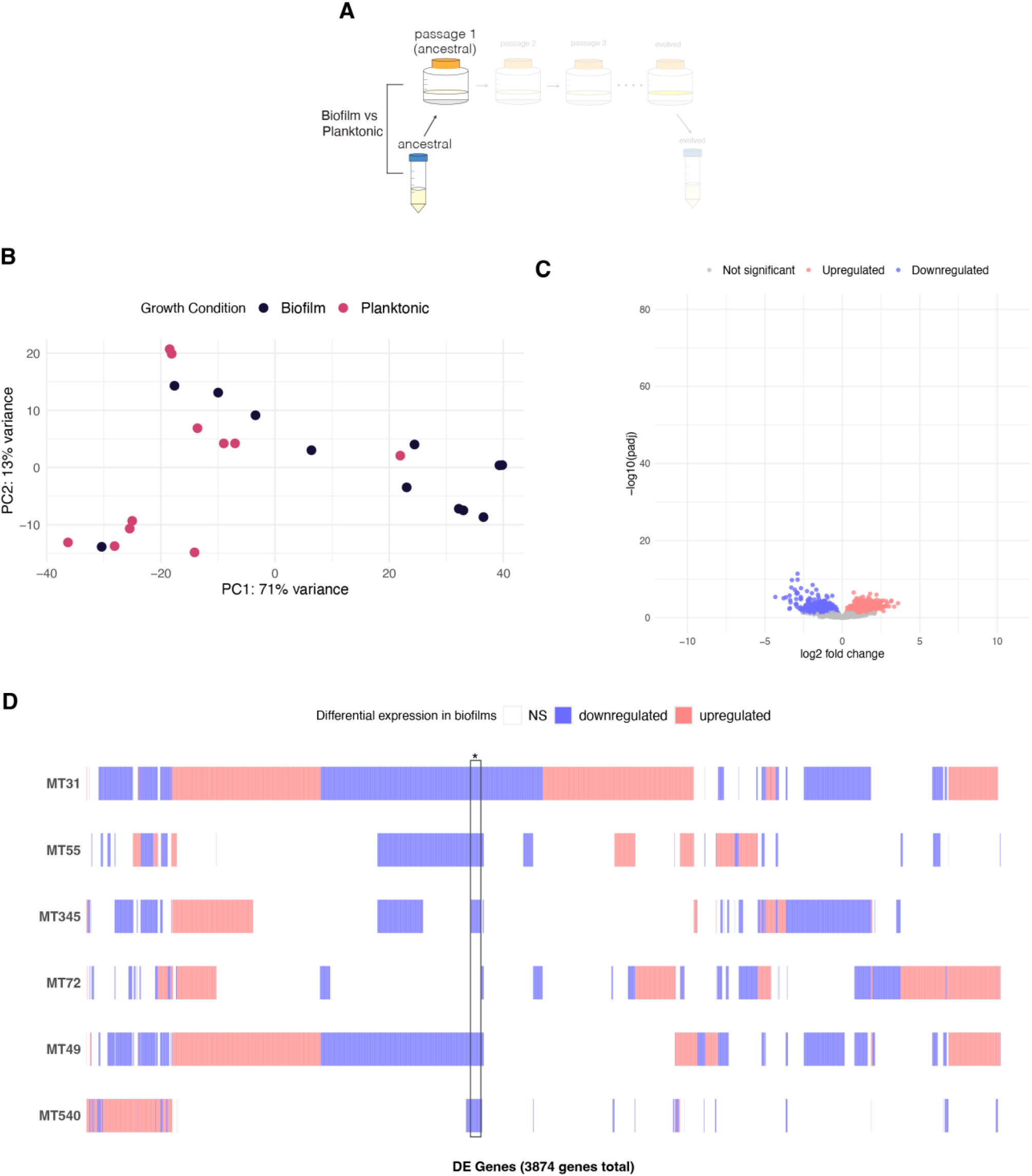
Genetic background affects transcriptional mechanisms of biofilm growth. A) Experimental diagram highlighting comparator populations: ancestral populations grown as pellicle biofilms are compared to the same populations grown in planktonic cultures. B) Principal component analysis (PCA) of normalized, log-transformed gene expression for ancestral populations. Each point is a biological replicate. Results demonstrate diversity among ancestral transcriptomes, with intermingling of planktonic and biofilm samples. C) Volcano plot summarizing differential expression between ancestral populations grown as biofilms and as planktonic cultures. Log transformed adjusted p-values plotted against the log2 fold change for each gene. Genes that did not have significant differential expression are shown in grey. Very few genes are significantly differentially expressed when all populations are analyzed together. D) Matrix of individual DEGs shared across ancestral populations. A total of 3874 DEGs are plotted according to whether the gene is upregulated (red), downregulated (blue) or not significantly differentially expressed (white) in each population. *Only 4% of downregulated genes are shared by at least 5 populations while no upregulated genes are shared across more than 4 populations.

### Biofilm passaging drives uniformity of the biofilm transcriptome and widespread repression of gene expression

Prior to biofilm passaging, we observed substantial diversity among biofilm transcriptomes in our sample (Figure 2, 3). The application of a uniform selection pressure resulted in a more uniform biofilm transcriptome across genetic backgrounds, with clear separation of samples from biofilm and planktonic conditions: approximately 76% of variance in our samples can be attributed to growth condition (represented by PC1, Figure 4B). When all populations were analyzed together, approximately 75% of the genome was involved in the transcriptomic response to biofilm conditions, with ∼ 3600 genes differentially expressed in comparison with planktonic growth (Figure 4C, Supplementary Data 2). In comparison with ancestral populations, the evolved biofilm transcriptome was characterized by even more extensive repression of gene expression (Figure 3C, 4C). Looking at the expression of individual genes, it is obvious that gene expression in evolved populations is more homogenous than in ancestral populations (Figure S2). This greater uniformity is also reflected in a larger proportion of shared, downregulated DEGs across populations: 17% in the evolved versus 4% in the ancestral populations (Figure 3D, 4D). We also observed a small proportion (6%) of DEGs that were upregulated across 5 populations, suggesting uniform recruitment of a cadre of genes under laboratory conditions (Figure 4D).

**Figure 4:**
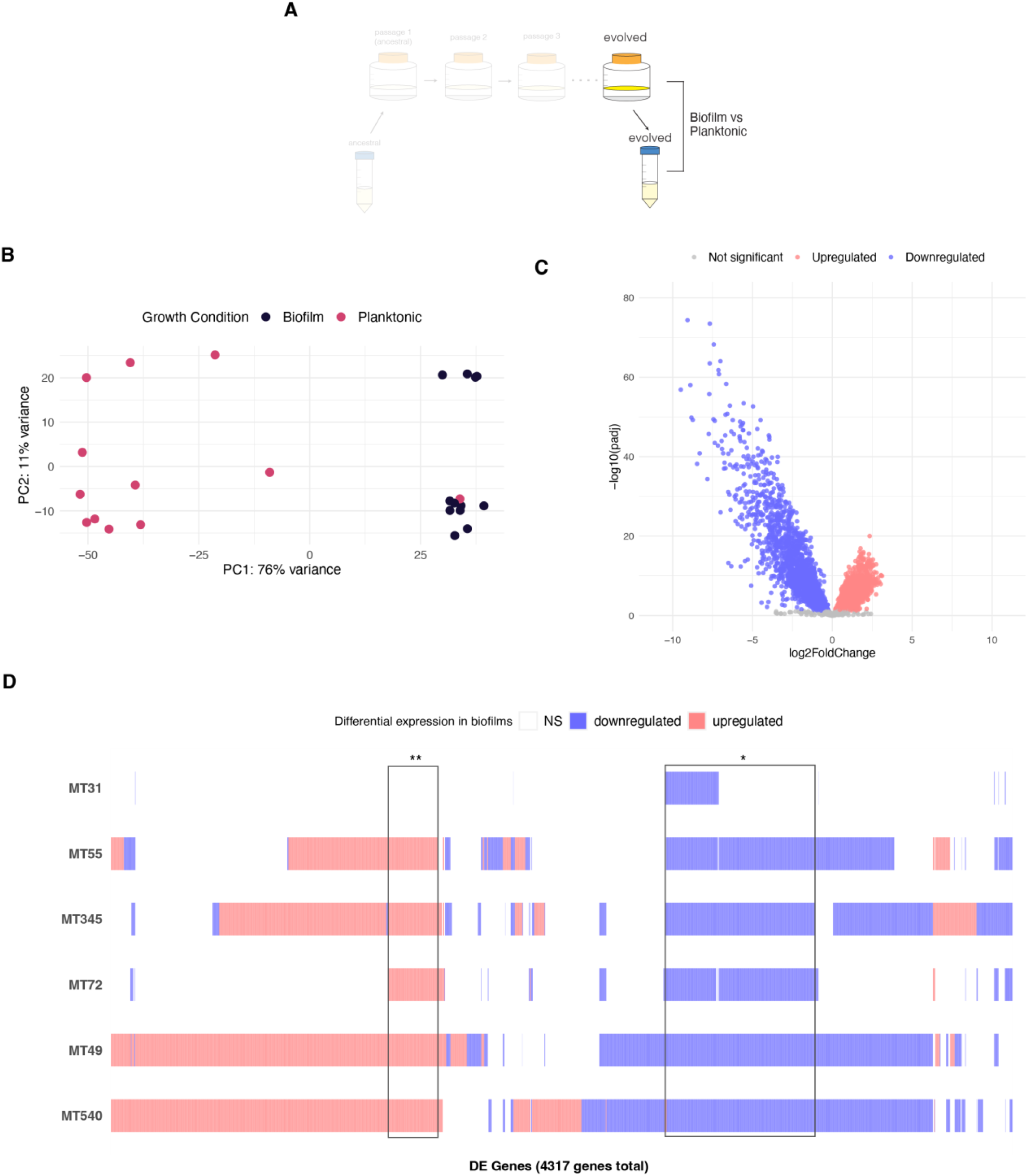
Transcriptional responses to biofilm growth following biofilm passaging. A) Experimental diagram highlighting comparator populations: evolved populations grown as pellicle biofilms are compared to the same populations grown in planktonic cultures. Analyses are parallel to those of ancestral populations in Figure 3. B) Principal component analysis (PCA) of normalized, log-transformed gene expression for evolved populations grown as biofilms and planktonic cultures. Biofilm passaging resulted in clear separation of biofilm and planktonic transcriptomes. These transcriptomes were not clearly separated at baseline (Figure 3B) C) Volcano plot summarizing differential expression between evolved populations grown as biofilms and as planktonic cultures. Log transformed adjusted p-values plotted against the log2 fold change for each gene. Genes that did not have significant differential expression are shown in grey. Biofilm transcriptomes were characterized by broad-scale downregulation of gene expression. D) Matrix of individual DEGs shared across evolved populations. A total of 3568 DEGs are plotted according to whether that gene is upregulated (red), downregulated (blue) or not significantly differentially expressed (white) in each population. 17% of downregulated genes are shared by at least 5 populations (*), while 6% of upregulated genes are shared by at least 5 populations (**). This contrasts with ancestral populations, which did not share any upregulated genes and only shared 4% of downregulated genes across 5 or more populations (Figure 3D).

Notably, there is one sample that is an outlier with respect to DEGs as well as its overall gene expression profile. This sample, obtained from the biofilm-passaged population of MT31 grown under planktonic conditions, has a gene expression profile more similar to a biofilm than to other planktonic samples (Figure 4B, S2B). If we classify this sample as planktonic, we identify few DEGs identified within this population (Figure 4D). If we instead group planktonic MT31 with the biofilm samples, we see a stronger contrast between conditions, and a much higher number of DEGs for MT31, many of which are shared with the other biofilm populations (Figure S3). After extended passaging as a biofilm, we attempted to grow our populations planktonically again and while most populations reverted to their ancestral planktonic transcriptomes, this replicate did not. Interestingly, this population was not defective for planktonic growth (as measured by OD600) and in fact evolved a higher rate of planktonic growth in response to biofilm passaging (*17*). One feature of these clinical strains is that unlike H37Rv they are not well adapted to laboratory growth and often develop biofilm-like “clumps” in planktonic culture, even in the presence of detergents. This phenomenon may explain why this particular replicate had a biofilm-like transcriptome, despite being grown in planktonic culture. We hypothesize that MT31’s transcriptome failed to shift to a planktonic pattern because it lost transcriptomic flexibility as a result of biofilm passaging, and/or that such flexibility was unnecessary because the biofilm transcriptome did not impose any fitness costs under planktonic growth conditions.

### Genetic background shapes adaptive trajectories

To further investigate how patterns of biofilm gene expression changed after passaging, we compared transcriptomes of *M. tb* passaged under selection for biofilm growth with corresponding ancestral populations (Figure 5). These analyses revealed three striking patterns. First, passaging under selection for biofilm growth reduced overall transcriptome diversity as summarized in a PCA (Figure 5A). Second, convergence of transcriptomes occurred in a lineage-specific pattern, with the evolution of clearly differentiated groups that corresponded with sub-lineage designations (Figure 5A). Finally, we found this pattern to be specific to biofilm growth as the transcriptomes of evolved populations grown planktonically remained intermingled and diverse (Figure 5B). These results indicate that transcriptomic responses were specific to the selection pressure applied, and that adaptation to this new environment was constrained by bacterial genetic background.

**Figure 5:**
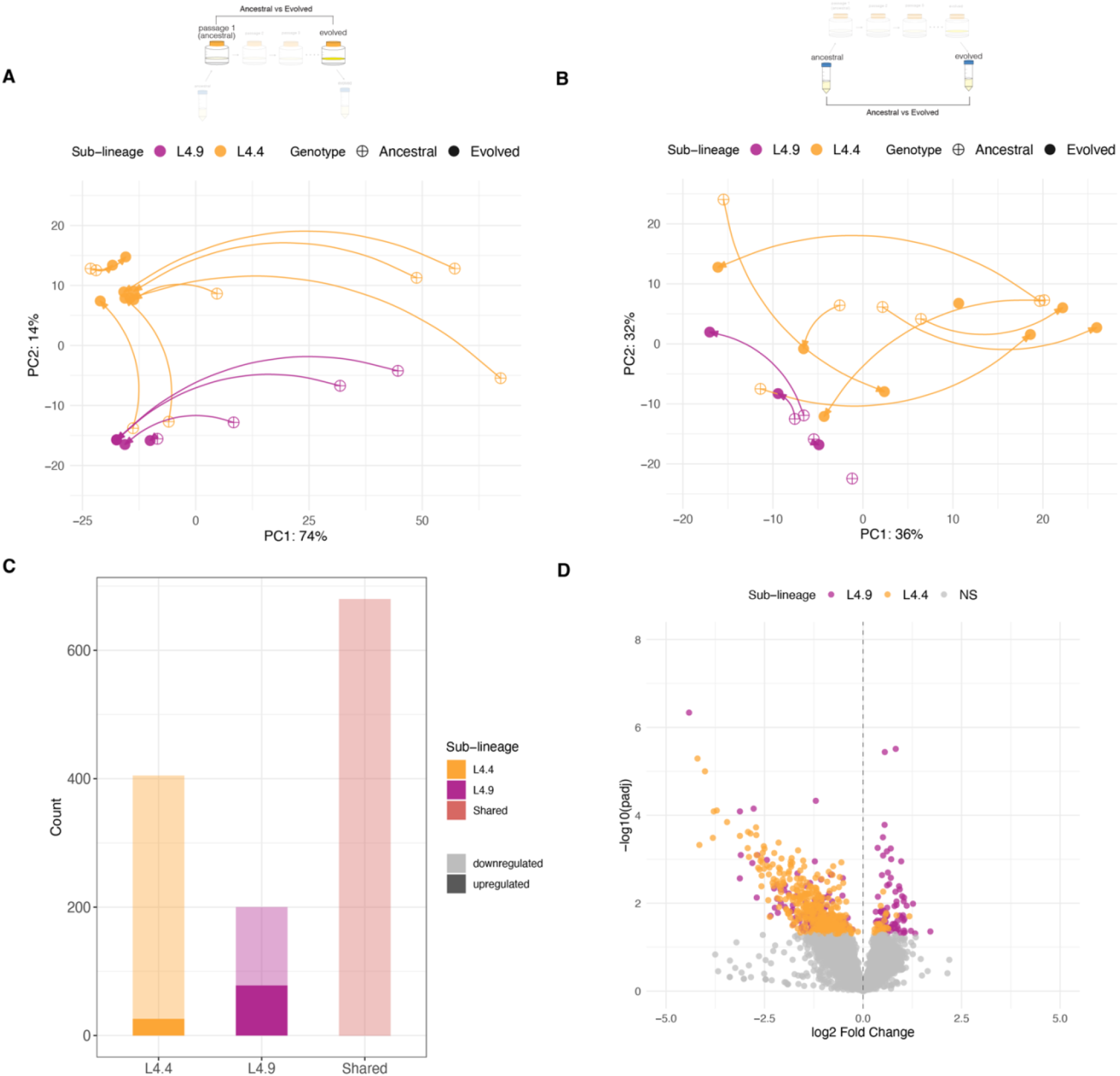
Genetic background shapes adaptive trajectories. A) Principal component analysis (PCA) of normalized, log-transformed expression counts from ancestral and evolved biofilm populations. Arrows are drawn between corresponding ancestral and evolved populations, indicating the trajectory of evolution across passaging. Points are colored by sub-lineage of the ancestral population. B) PCA of normalized, log-transformed expression counts from ancestral and evolved populations grown under planktonic conditions. Like panel A, arrows are drawn between corresponding ancestral and evolved populations and are colored by sub-lineage. Biofilm passaging reduced transcriptome diversity, with populations converging on a sub-lineage-specific signature (A); this pattern was not observed under planktonic growth conditions (B). C) Bar plot of DEGs comparing evolved and ancestral biofilm populations broken down by sub-lineage. There are 680 DEGs shared between sub-lineages, and 405 and 200 DEGs unique to L4.4 and L4.9, respectively. Opacity of the bar indicates the subset of DEGs that are either up (dark) or downregulated (light) after passaging. The pellicle biofilm transcriptome is characterized by broad scale downregulation of gene expression with a smaller complement of genes that are upregulated in a sub-lineage specific manner. D) Differential expression of DEGs unique to each sub-lineage, from the comparison of evolved to ancestral biofilm populations. Log transformed adjusted p-values plotted against the log2 fold change for each gene. Genes that did not have significant differential expression are shown in grey. Lineage 4.9 evolved a larger complement of upregulated genes in response to biofilm passaging.

Gene-by-gene analyses of the transcriptome also revealed lineage-specific patterns. As noted above, biofilm growth was associated with widespread downregulation of gene expression (compared to exponential growth in planktonic culture). Biofilm passaging reinforced this pattern, with comparisons of evolved and ancestral populations revealing further genome-wide downregulation (Figures 5C, 5D). All differentially expressed genes (DEGs) that were shared among sub-lineages were downregulated (Figure 5C). Outside of these shared DEGs, the sub-lineages exhibited distinct patterns with sub-lineage 4.9 showing a larger proportion (39%) of upregulated genes in its DEG repertoire than sub-lineage 4.4 (6%) (Figure 5C, 5D, Supplementary Data 3). We found previously that *M. tb* populations within sub-lineage 4.9 all evolved large duplications (*17*), raising the possibility that the relatively large proportion of upregulated genes observed here arose from the duplication. However, the majority (55%) of L4.9-specific, upregulated DEGs lie outside the duplicated region (Supplementary Data 3). Thus, the increased proportion of upregulated genes in L4.9 does not appear to arise directly from increased gene dosage from the genomic duplication. It is also noteworthy that while all populations in L4.4 evolved similar transcriptomes (Figure 5A), we did not identify any genomic mutations that were shared across these populations (*17*). Together, these observations implicate a complex relationship between genetic backgrounds, novel mutations, transcriptional responses, and phenotypes. It appears that the application of a uniform selection pressure can result in similar phenotypic adaptation (in this case enhanced biofilm growth) via distinct mutational paths, that result in similar transcriptional responses. Thus, when exposed to a new environment, sub-populations of *M. tb* may adapt similarly using different paths to the same goal.

### Expression consequences of large tandem duplication and intergenic SNP

In our passaging experiment, a near identical genomic duplication emerged repeatedly and specifically in association with biofilm passage, providing powerful evidence for its association with biofilm growth (*17*). We have termed this the MMMC duplication, an umbrella term for duplications starting around 3.1Mb (*Rv3100)* with lengths from 150-350 kb, which have been described before both *in vitro* (*20*–*23*), and *in vivo* (*24*). In our experiments, the independently arising MMMC duplications became fixed in biofilm passaged populations and were stable for up to 20 passages, whereas they were absent from populations passaged as planktonic cultures (*17*). Here we sought to investigate the impact of a large-scale duplication on the biofilm transcriptome, thereby illuminating potential mechanisms for its impact on biofilm growth. A more in-depth analysis of the coordinates of the MMMC duplications using sliding-window DNA sequencing coverage revealed them to be larger than we had originally reported (∼175 versus 120 kb) using another method of structural variant identification (*17*). The duplication has slightly different coordinates in each population, but spans from position ∼3.54 to 3.76 Mb, with an average length of 175 kb, and includes genes *hpx* through *PPE56* (Figure 6B).

**Figure 6:**
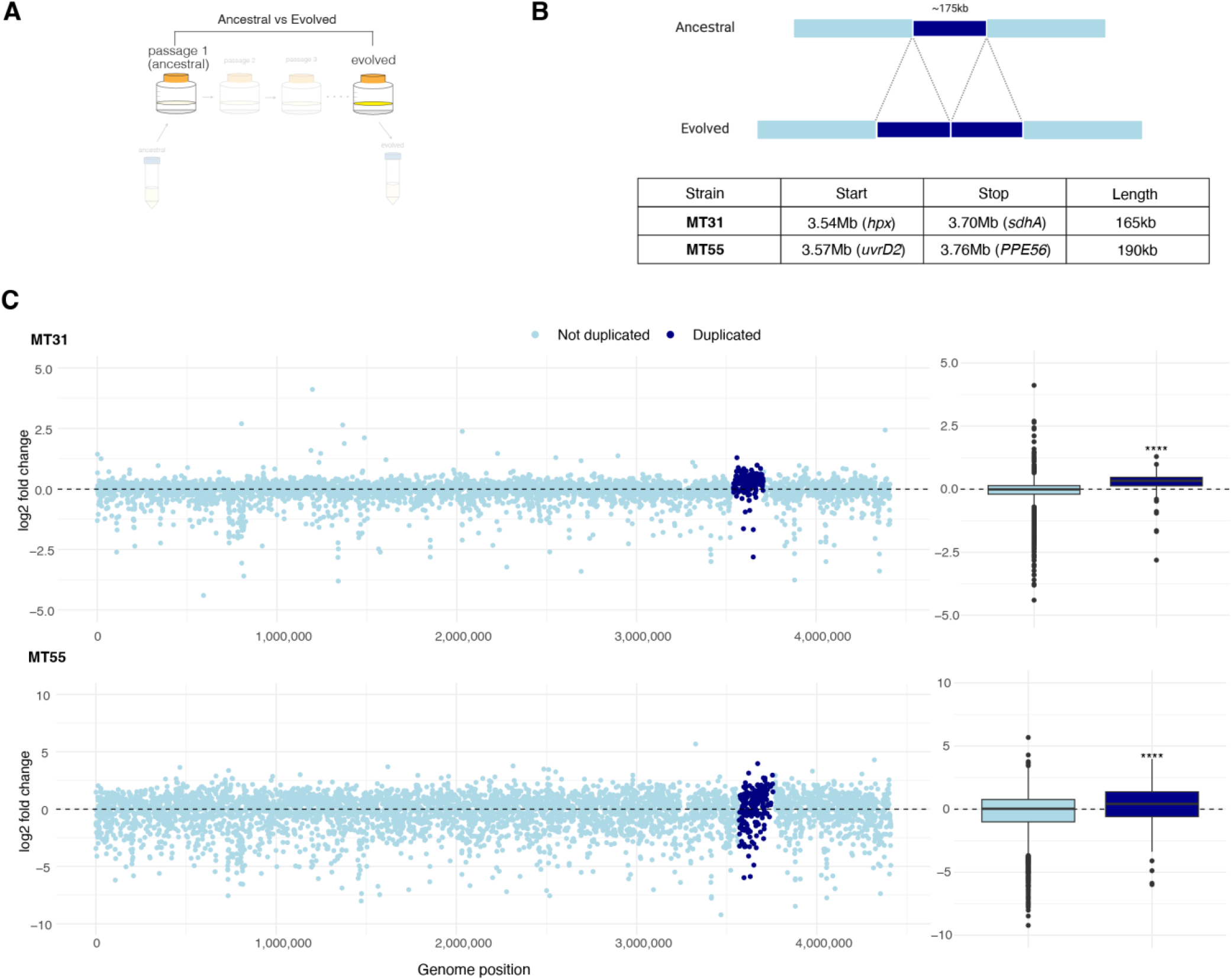
Genome duplication results in complex patterns of gene expression. A) Experimental diagram highlighting comparator populations: comparison of transcriptomes between evolved and ancestral biofilm populations. B) Top: large tandem duplication that evolved in populations MT31 and MT55 under selection to grow as a biofilm. Bottom: coordinates of the duplication vary slightly between populations, with an average duplicated length of 175 kb. C) Left: log2 fold changes in gene expression between evolved and ancestral biofilm populations plotted against the position in the genome. Each point is a single gene, colored according to whether that gene lies inside of the duplicated region for that population. Right: Boxplots of log2 fold changes in gene expression, for genes outside (light blue) and inside (dark blue) the duplication. Genes within the duplication were significantly more upregulated in both populations (Mann Whitney U test with Benjamini-Hochberg correction, *p* < 0.0001).

A priori, we might expect that a second copy of a given gene would increase expression by 2x, corresponding to a log2 fold change (L2FC) value of 1. Such a modest fold change is not likely to be identified as statistically significant and in fact only 39 duplicated genes were significantly upregulated in either MT31 or MT55 (Supplementary Data 3). To further investigate impacts of the MMMC duplication on gene expression, we plotted L2FC values between evolved and ancestral biofilm populations (Figure 6A) for all genes across the genome and compared changes in duplicated genes to non-duplicated genes (Figure 6C). Overall patterns of L2FC values seem to differ between the two populations: MT55 has greater differentiation between ancestral and evolved populations, resulting in L2FC values of higher magnitude than MT31 (Figure 6C). The distribution of L2FC values within the duplication is shifted upward relative to genes outside the duplication, a difference that was statistically significant under both biofilm and planktonic growth conditions (Mann Whitney U test with Benjamini-Hochberg correction, *p* < 0.0001) for both MT31 and MT55 (Figure 6C, Figure S4B). However, we did not observe a simple two-fold increase in gene expression within the duplication: genes exhibited a range of values with some strongly down- and up-regulated. Thus, the transcriptomic impacts of the gene duplication appear to be complex, rather than being simply characterized by a doubling of gene dosage.

We expected expression changes of genes within the duplication to be specific to evolved populations of MT31 and MT55. However, we found two other populations with expression changes within this region: expression was overall downregulated in evolved populations of MT49 and MT72 under biofilm conditions (Figure S4A) and in MT49 under planktonic conditions (Figure S4B). The difference was relatively subtle for MT72 but striking for MT49 (Figure S5A). Coverage plots from DNA sequencing reads did not reveal a genomic deletion in this region for MT49 (Figure S5B). Using coverage of RNA sequencing in this region we observed that the ancestral population had relatively high expression of genes in this region compared to the surrounding area (Figure S5B). Thus, it appears that the difference in gene expression at this locus arises from increased expression in the ancestral population, rather than decreased expression in the evolved population. Our hypothesis is that a transient MMMC duplication arose in the ancestral population during the process of growing the population as a biofilm for RNA extraction and sequencing. This is further supported by RNA coverage of ∼1.25x within this region indicating that the duplication was present at intermediate frequency when the sample was sequenced (Figure S5B).

Our observation that duplications in this region of the *M. tb* genome appear repeatedly in association with a selective pressure to grow as a biofilm (*17*) suggest that genes within the duplication may be important for robust biofilm formation. Our hypothesis that this adaptation affects fitness via gene dosage appears to be supported by the transcriptomic data, although the relationship between duplicated genes and increased expression is complex. To our knowledge, these data represent the first in-depth description of the effects of large-scale genome duplications on gene expression in bacteria, expanding our understanding of these variants in the broader context of strategies for bacterial adaptation to new environments.

In another instance of convergent adaptation, L4.4.1.1 strains MT49 and MT540 both acquired intergenic SNPs upstream of *lpdA* (Figure S6A). We previously showed using quantitative PCR (qPCR) that these intergenic SNPs led to increased expression of the downstream gene (*lpdA*), likely by interfering with a known transcription factor binding site in this intergenic region (*17*). Here we used the transcriptomics data to look at the rest of the genes in the operon downstream of *lpdA*, a total of 6 genes. We found that that the intergenic SNPs had variable effects on expression of downstream genes under biofilm conditions with some genes upregulated (*lpdA, glpD2, Rv3300c* and *lpqC*) while others were downregulated (*phoY1* and *atsB*), indicating that these genes are not all co-transcribed (Figure S6). Because this operon lies within the bounds of the MMMC duplication the analysis of MT49 under biofilm conditions was also confounded by the transient duplication as we described above for the MMMC duplication (Figure S6). Like the duplication these SNPs appear in association with biofilm growth and appear to have complex effects on gene expression.

### Non-coding RNA and small proteins are a part of the adaptive transcriptome

Advances in transcriptomic and proteomic methods have revealed an abundance of non-coding RNA (ncRNA) (*25*–*31*) and small open reading frames (sORFs) (Shell et al., 2015; C. Smith et al., 2022) in the *M. tb* genome. In *M. tb*, ncRNA are important for gene regulation and adaptation (*34*) and virulence (*35*). In other bacteria, diversity in the non-coding transcriptome is hypothesized to be a driver of inter-strain divergence and adaptation (*36*). We hypothesize that these features may similarly play a key role in *M. tb* adaptation to biofilm growth. We included a custom list of ncRNA and sORFs (Supplementary Data 4) in our annotation file used for all differential expression analyses in Figures 2-6. To observe subtle patterns in expression of these non-canonical features we separated them from the rest of the open reading frames (ORFs) and looked at their differential expression across biofilm passaging, and in the evolved populations.

We observed that in general, sORFs and ncRNA follow similar patterns to coding regions: there is more downregulation (both in the number of the features and the magnitude of differential expression) than upregulation (Figure 7A,B), and across passaging the expression of these features converge in a genetic background dependent manner (Figure 7C), a phenomenon that does not hold true under planktonic growth conditions (Figure S7). This reflects what we see with transcriptome-wide adaptation to biofilm growth, which is that specific, directional changes in gene expression as a result of the applied selective pressure (biofilm growth in this case), are only observed when that selective pressure is maintained (Figure 5A,B). One pattern that was particularly striking was the upregulation of ncRNAs in evolved biofilms: when all populations were analyzed together, ncRNA represent 33% of all strongly upregulated features (L2FC > 2, Supplementary Data 2) despite comprising only 12% of all features. Integrating the analysis of all populations with those of individual populations, we find 14 ncRNA that are strongly upregulated in at least 5 of our individual populations (Table 1). Comparing the distribution of log2 fold change values of ncRNA with other feature types (ORFs and sORFs), we found that ncRNAs were significantly more likely to be upregulated in ancestral biofilms (Figure 8A), as well as being significantly more likely to be upregulated as a result of passaging (Figure 8B). Our results demonstrate that upregulation of ncRNA seems to be a common feature of the *M. tb* biofilm transcriptome, and that non-canonical features such as small proteins and non-coding RNAs play an important role in bacterial adaptation to new environments.

**Figure 7:**
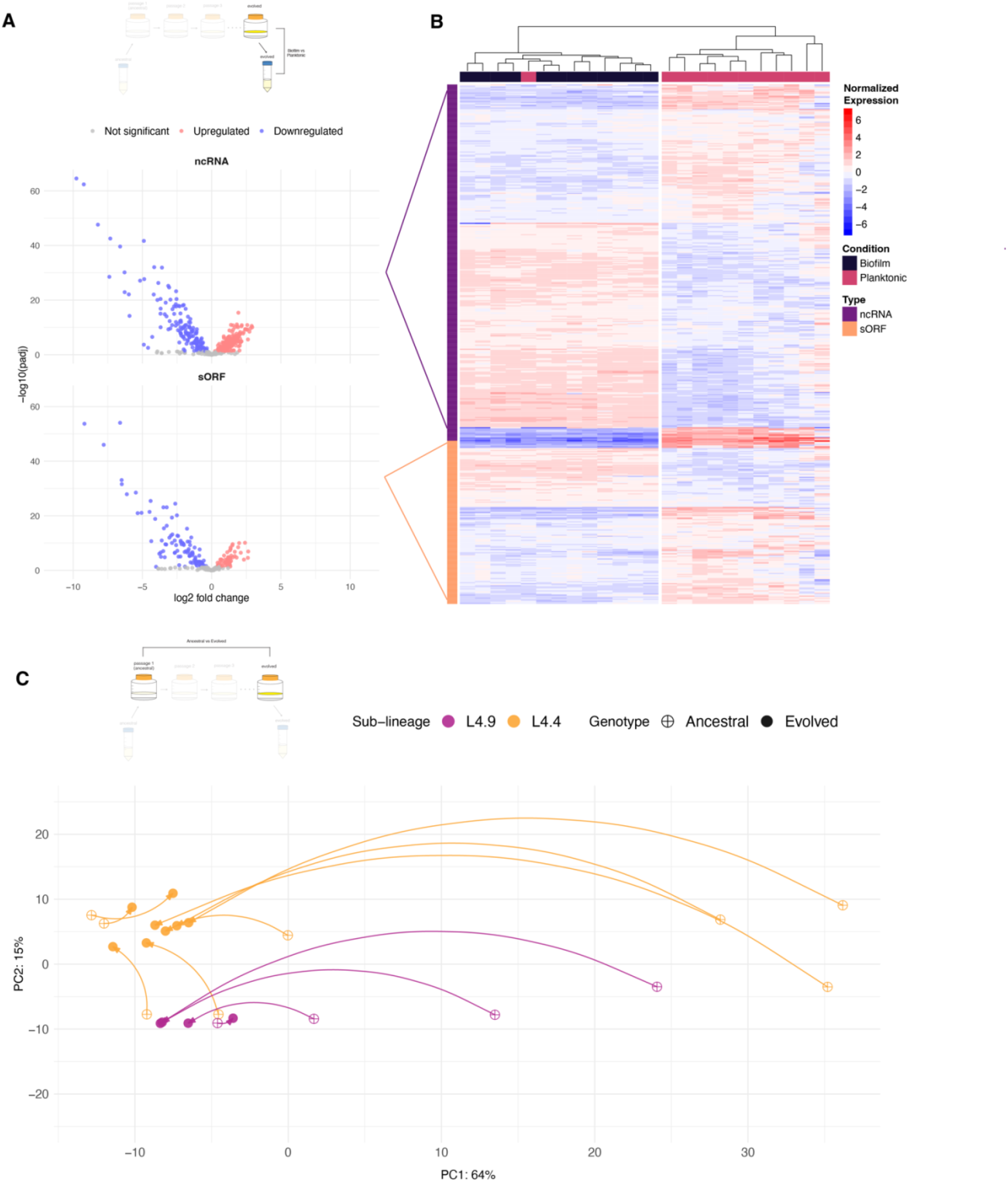
Expression of ncRNAs and sORFs mirror coding regions after passaging. A) Volcano plot summarizing differential expression of ncRNAs and sORFs in evolved populations. Log transformed adjusted p-values plotted against the log2 fold change for each gene. Genes that did not have significant differential expression are shown in grey. B) Heatmap of normalized, log-transformed expression counts for ncRNAs and sORFs with significant differential expression in evolved populations. Each column is a single sample from an evolved population, grown either as a biofilm or in a planktonic culture. Each row is a gene labeled either ncRNA or sORF. Expression values for each gene are normalized to the mean across samples. Samples are clustered by Euclidean distance and plotted as a tree at the top of the heatmap. C) Principal component analysis (PCA) of normalized, log-transformed expression counts of ncRNAs and sORFs from ancestral and evolved biofilm populations. Arrows are drawn between corresponding ancestral and evolved populations, indicating the trajectory of evolution across passaging. Points are colored by sub-lineage of the ancestral population. The effects of biofilm passage on ncRNAs and sORFs mirror that of coding regions: reduced transcriptome diversity and widespread downregulation with populations converging on a sub-lineage-specific signature.

**Figure 8:**
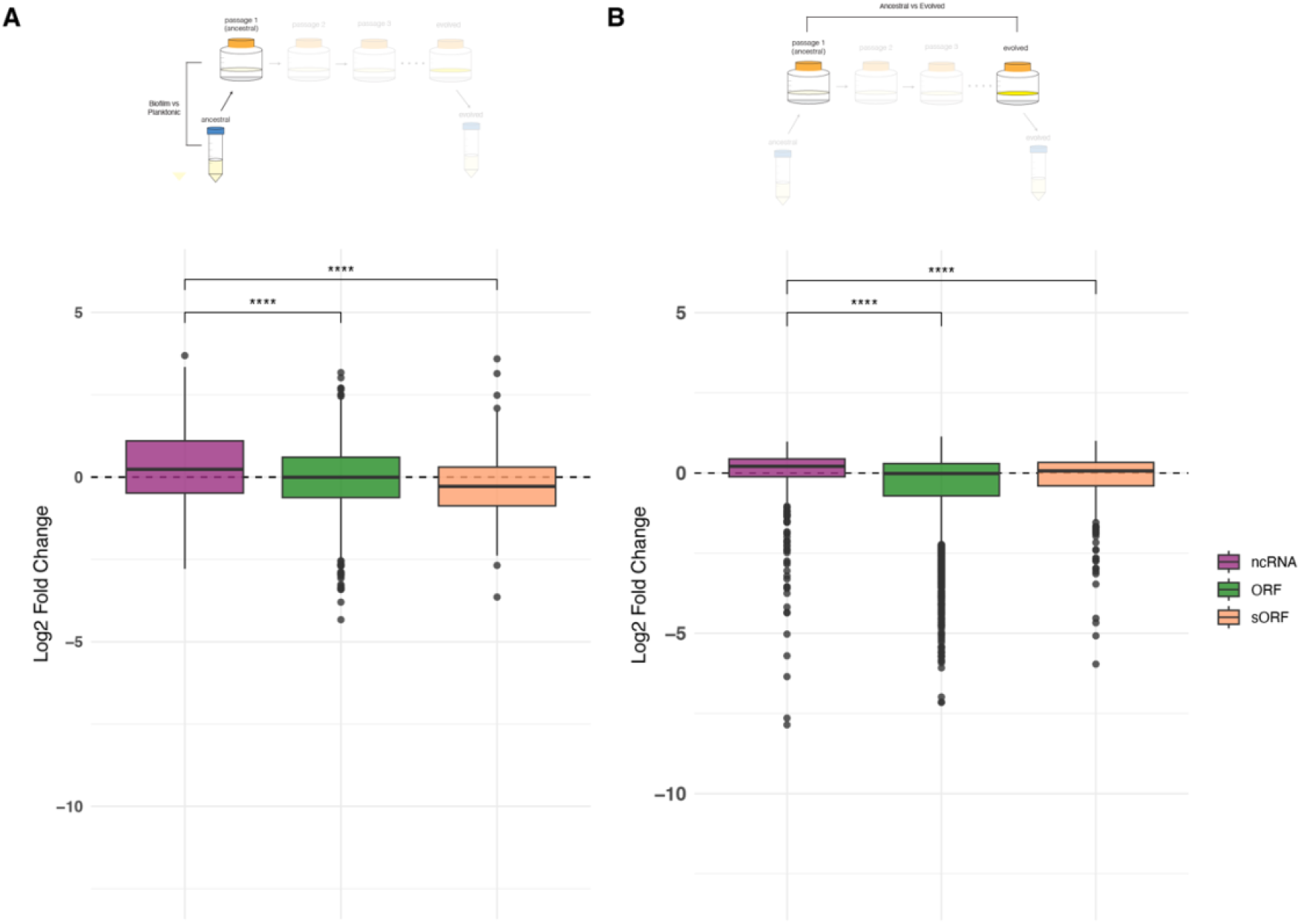
Upregulation of ncRNA is a common feature of *M. tb* biofilm growth. A) Log2 fold change (L2FC) values from ancestral populations by feature type. ncRNA have significantly higher L2FC values than ORFs and sORFs (Mann Whitney U Test with Benjamini-Hochberg correction, *p* < 0.0001) indicating they are more likely to be upregulated in biofilms. B) L2FC values between evolved and ancestral biofilms by feature type. ncRNA expression increased significantly more than either ORFs or sORFs after biofilm passage (Mann Whitney U Test with Benjamini-Hochberg correction, *p* < 0.0001).

**Table 1:**
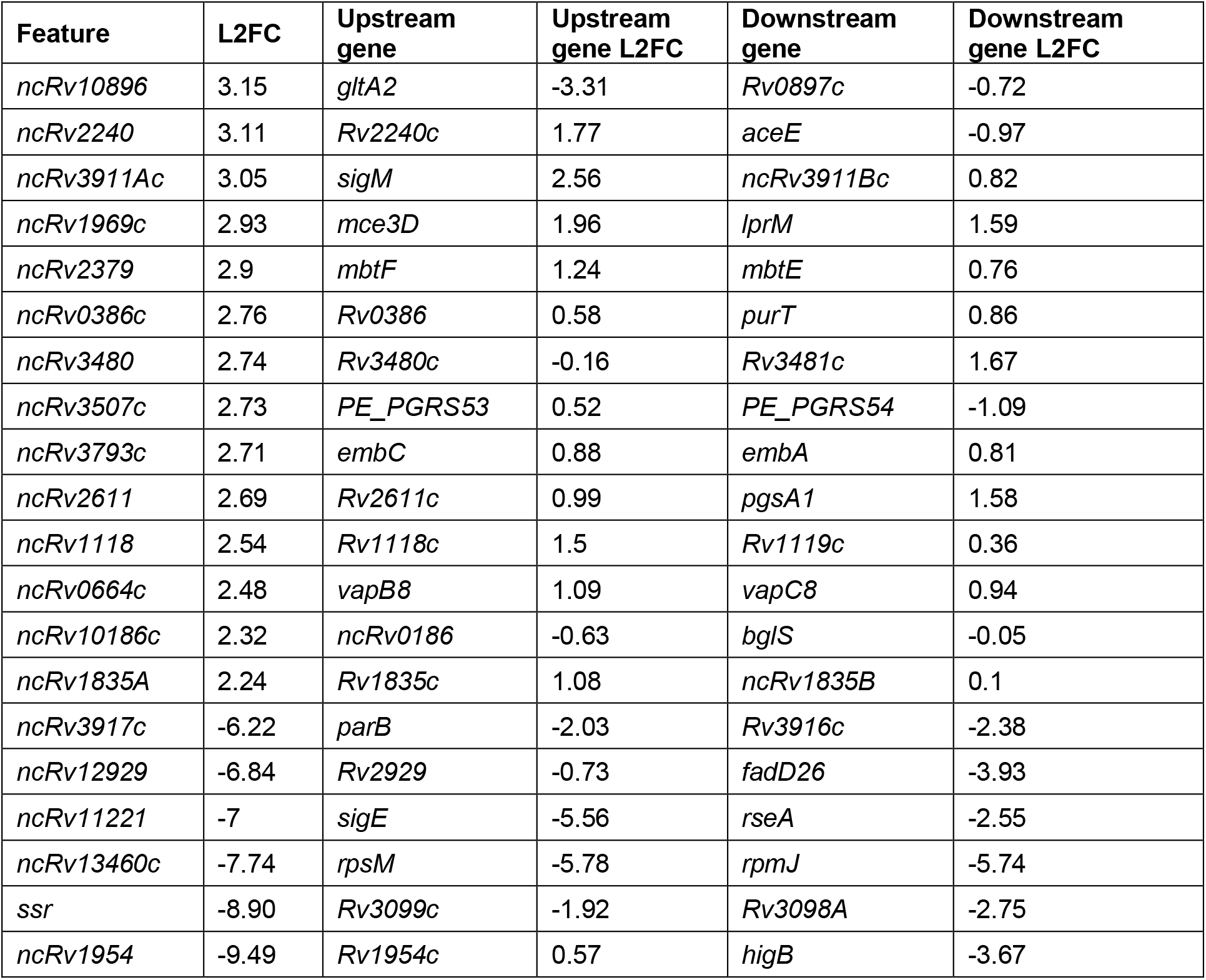
Most significantly downregulated (< -5 L2FC) and upregulated (> 2 L2FC) ncRNAs in evolved populations. Features included in this list are significantly differentially expressed when all populations were analyzed together, as well as in at least five individual populations. Log2 fold change (L2FC) values given for analysis of all populations together.

## Discussion

Here we have characterized the diversity of transcriptomes among closely related clinical strains of *M. tb*. We observed genome-wide changes in gene expression in response to the application of a specific selective pressure, resulting in convergence on a biofilm transcriptome that was structured by lineage. Characterization of the pellicle transcriptome revealed a complex relationship between gene dosage, gene expression and the upregulation of non-coding RNA. Overall, this work highlights the complexity of the regulatory system in *M. tb* and provides a framework for considering the transcriptome as a target of selection.

### M. tb biofilm transcriptome: one size does not fit all

Little is known about the structure and function of *M. tb* biofilms (*37*), and even less about the transcriptional signatures that define this growth form. To our knowledge there is just one prior study of pellicle biofilm transcriptomics (*16*), which pinpointed a role for isonitrile lipopeptide (INLP) in biofilm growth. In their study of an attenuated strain of *M. tuberculosis* (mc^2^7000), Richards et al. discovered that a 6-gene cluster termed the INLP synthase complex (*PPE1, Rv0097, fcoT, fadD10, Rv0100, nrp*) was highly upregulated during pellicle growth and they performed further experimental work to identify INLP in mycobacterial biofilms.

In the present study of closely related, fully virulent clinical strains of *M. tb*, we found that the pellicle biofilm transcriptome exhibited substantial variation among strains (Figures 2, 3). This variation is evident in the number of genes differentially expressed under biofilm growth conditions (Figure 2), as well as the identity of genes subject to differential expression. As a result, there are few differentially expressed genes that are shared among strains (Figure 3C, 3D) and strain-to-strain variation dwarfs transcriptomic signals differentiating planktonic and biofilm growth (Figure 3B). Among this variation, we did identify one strain that exhibited a similar pattern to the strain studied by Richards et al. Four out of the six genes in the INLP synthase complex were upregulated and in the 98th percentile of differentially expressed genes for the strain MT55 (Supplementary Data 1). Genes in the complex were not upregulated to any significant degree in any of the other strains in our study, indicating that INLP may be a significant component of the extracellular matrix in some, but not all strains. Integrating our results with those of Richards et al., we conclude that natural populations of *M. tb* exhibit substantial variation in the regulatory mechanisms underlying biofilm growth, such that there appear to be multiple paths to this complex phenotype.

### Flattening and global repression in evolved populations

The application of a uniform selection pressure resulted in evolution of a more uniform transcriptome, loss of strain-to-strain variation, and disambiguation of gene expression under conditions of biofilm growth (Figure 4). The evolved transcriptome is characterized by widespread repression of gene expression, a pattern that was evident in ancestral populations but markedly intensified after passaging (Figure 3C, 4C). Of the ∼3600 genes differentially expressed across all pellicle passaged populations, there are similar numbers of down and upregulated genes (Figure 4C), however, the magnitude of the most downregulated genes exceeded 3x the magnitude of the most highly upregulated genes (Supplementary Data 2). Similar patterns of widespread repression have been identified previously, in studies of bacteria isolated from sputum (*38*, *39*) and in vitro models of dormancy (*5*). Where the pellicle model differs from sputum and dormancy, however, is in the global regulatory schemes that dictate the transcriptional program. The DosR regulon controls a group of genes that are activated under anaerobic conditions to trigger entry into a non-replicating dormant state (*40*). Expression of the DosR regulon is a feature of the dormancy transcriptome (*5*) and has also been shown to characterize the sputum transcriptome (*38*, *41*, *42*). Data from Richards et al. show that the DosR regulon was largely downregulated in their pellicle model, relative to exponential growth (*16*). Here, we found that except for one strain (ancestral MT72), all ancestral and evolved biofilms showed significant downregulation of DosR regulon genes (Supplementary Data 1,2). Thus, it appears that while gene expression during pellicle growth shares some similarities to sputum and dormancy, the mechanisms of adaptation to pellicle growth are strain variable, and appear to occur most commonly by a distinct, DosR-independent mechanism.

### Signatures of pellicle adaptation in the non-coding transcriptome

Growth as a biofilm was evident in the non-coding transcriptome, in addition to the coding transcriptome. Expression of non-coding RNAs (ncRNAs) was increased relative to that of genes during biofilm growth of ancestral populations (Figure 8A). Evolution under selective pressure for biofilm growth reinforced this pattern with ncRNA being significantly more upregulated than other loci after passaging (Figure 8B). Considering both ncRNAs and small open reading frames (sORFs), adaptation to pellicle growth resulted in emergence of a uniform biofilm transcriptome that was clearly distinguishable from planktonic conditions (Figure 7). We identified 14 ncRNAs that were significantly upregulated (L2FC > 2) and 6 that were significantly downregulated (L2FC < -5) in our pellicle passaged populations (Table 1). These are candidate loci mediating shifts in gene expression that enable *M. tb* growth as a biofilm. It has been shown that upregulation of even a single ncRNA can induce widespread gene repression (*25*, *28*). There are many targets of ncRNA regulation including genes involved in lipid metabolism (*26*), response to iron-limitation (*28*) and two-component regulatory systems (2CRS) (*30*). Therefore, it is hypothesized that in *M. tb,* ncRNA mediate a rapid response to new environmental stimuli, whereby changes in expression of a small number of ncRNA can have large impacts by regulation of many other genes. A similar phenomenon was recently uncovered in *C. difficile* where ncRNAs were shown to control initiation of sporulation (*43*). We show here that ncRNAs are upregulated in our pellicle biofilm model and hypothesize that upregulation of ncRNAs contributes to the widespread changes in gene expression (particularly downregulation) we see in pellicle passaged populations, a phenomenon we have termed Non-Coding Regulatory Reprogramming (NCRR). Future work isolating the downstream effects of individual ncRNA on expression may further illuminate the role of NCRR in the complex regulatory network of *M. tb*.

### How do transcriptomes evolve?

Experimental evolution studies have shown that many adaptive mutations in bacteria are predicted to have a regulatory effect. Studies of *Escherichia coli* (*44*–*46*) and *Pseudomonas aeruginosa* (*47*) indicate that rapid adaptation to new environments can be accomplished with a few well-placed mutations in regulatory genes, a pattern we observed in our experiment evolving *M. tb* (*17*). What is less well-known however, is the impact of these mutations on the transcriptional landscape. We sought to characterize the effects of these adaptive mutations on the transcriptome and found the most obvious pattern to emerge after passaging was the intensification of pre-existing patterns observed in our ancestral populations. Selection for pellicle growth resulted in stronger downregulation of genes (Figure 3C, 4C), reduction of sample-to-sample diversity (Figure 3B, 4B) and further upregulation of ncRNA (Figure 8B). Of particular interest is the reduction of diversity between biofilm transcriptomes after passaging, an indication that these populations converged onto a shared transcriptome under the same selective pressure. These findings extend observations from previous studies illustrating parallel transcriptome adaptation after imposition of a uniform selection pressure. Under laboratory conditions, convergence towards a shared genic transcriptome has been observed several times in bacteria (*47*– *50*) and even in fungus (*51*). One study of *P. aeruginosa* transcriptomes from an explanted lung showed that genetically distinct strains exhibited minimal variability in their transcriptomes (*52*), indicating that this phenomenon likely holds true for bacterial adaptation during infection. Here, we have shown that the phenomenon applies to *M. tb*, and that it also extends to the non-coding transcriptome.

We identified a striking phenomenon beyond this convergence on a shared transcriptome. Our findings suggest that the adaptive trajectories of both the coding and non-coding transcriptomes are shaped by the genetic background of the bacterial population (Figures 5A, 7B). Populations with differentiated transcriptomes at baseline evolved shared, lineage-specific transcriptomes under selection for biofilm growth. This pattern was specific to biofilm growth, as neither convergence nor an effect of genetic background were evident during planktonic growth of evolved populations (Figures 5B, S7). There are several, not mutually exclusive, explanations of these phenomena. In our prior study describing genomic adaptations during this experiment, we found evidence that strain genetic background affects the types of mutations observed during selection for pellicle growth (*17*). Convergence on a strain-specific transcriptome could thus reflect the mutational trajectory associated with each genetic background, since the majority of observed mutations appeared to act by a regulatory mechanism (*17*). We discuss this possibility in more detail below.

### Transcriptomic consequences of mutations

One of the mutations that exhibited an association with lineage is a 175 kb duplication we termed the MMMC duplication. The MMMC duplication has been observed previously, and structural variants appear to be common in *M. tb* populations, likely facilitated by recombination among IS elements (*22*). We previously found the MMMC duplication to be specifically associated with pellicle adaptation and the L4.9 background. Here, we found that *M. tb* populations from the L4.9 background, known to have developed the MMMC duplication, converged on a similar transcriptome (Figure 5A).

The evolutionary history of *M. tb* is littered with evidence of structural variants, particularly duplications, which have formed some of the most important gene families like the Type VII secretion systems (*53*) and PE/PPE genes (*54*). In addition to the creation of new gene functions, duplications are also known to increase fitness through gene dosage effects. For example, an attenuated strain of *M. tb* missing ESX-3 effectors *esxG* and *esxH* developed duplications of a set of paralogous effectors which, when overexpressed as a result of duplication, compensate for the loss of *esxG* and *esxH* (*55*). Despite their prevalence in *M. tb*, transcriptional impacts of large structural variants are not well understood. A prior study of a genomic deletion found that it was associated with loss of expression of genes within the deletion (*56*), but to our knowledge there are no prior studies investigating transcriptional impacts of large duplications. Here we sought to identify the transcriptional impacts of the MMMC duplication. We identified a two-fold increase in gene dosage from the MMMC (*17*), thus we might naively expect the duplication to result in a two-fold increase in expression of genes within the duplication. However, this was not the case. Effects of the duplication on gene expression appeared to be highly variable, with some genes exhibiting expected two-fold increases (log2 fold change of 1) and others with as much as 9-fold increases (log2 fold change of ∼3) and 50-fold decreases (log2 fold change of ∼ -5) (Figure 6C, Supplementary Data 3). These findings illuminate a complex relationship between mutation, gene expression, and phenotype that belies a simple one to one map between gene dose and gene expression. Our initial hypothesis for the mechanism underlying this complex pattern is that the duplication of such a large region in the genome interrupts the function of trans-regulatory elements that are either encoded within, or act on loci within the duplication. Duplication of binding sites for a trans-regulatory element outside the duplicated region would effectively ‘dilute’ the effect of that regulator. Conversely, increased gene dosage of regulators within the duplication may affect expression of genes within and outside the duplication. In these scenarios the compounded effects of increased gene dosage and complex regulatory architecture could lead to highly unpredictable expression of genes in a large-scale duplication. Further analysis of the specific regulators and regulatory binding sites within the MMMC duplication may help to elucidate patterns in the expression of duplicated genes.

Another possible explanation for the apparently complex relationship observed between gene expression and gene dosage is modification of the duplication’s impact by concomitant mutations. However, the data do not support this hypothesis. The two strains from L4.9 did not share any mutations other than MMMC (*17*), and yet they evolved similar transcriptomes (Figure 5A). In fact, populations within L4.4 also converged on similar coding and non-coding transcriptomes (Figure 5A, 7B), and while L4.4.1.1 strains MT49 and MT540 share intergenic SNPs (Figure S6), there are no shared mutations across all L4.4 strains (*17*). Thus, convergence of the transcriptome does not appear to be driven by mutational convergence: different mutations on similar genetic backgrounds can result in similar transcriptomes. Transcriptome diversity has been described previously among clinical isolates of *M. tb* (*57*–*59*). Our observations here suggest that this diversity reflects both canalization (the tendency of a population to produce the same phenotype regardless of genotype) arising from genetic structure, including neutral haplotype and lineage structure, as well as variation in selection pressures encountered during natural infection.

### Implications & future directions

We know that biofilms represent heterogeneous populations, with sub-populations that differ in genotype and functional specialization (*60*). Recent work using spatial transcriptomics has also begun to reveal the ways in which bacterial communities coordinate gene expression to form biofilms (*61*, *62*). Our analyses in this work have compressed the diversity of the *M. tb* biofilm transcriptome within this population to a single dimension. Future work studying the diversity of gene expression within biofilm sub-populations will give us more insight into the form and function of *M. tb* biofilms.

In this study we have characterized both the patterns of gene expression within *M. tb* biofilms, as well as how those patterns change upon the application of a complex selective pressure. We discovered a cadre of upregulated ncRNA that we believe to be involved in genome-wide gene expression modulation, a phenomenon we call Non-Coding Regulatory Reprogramming (NCRR). We have further illustrated the complex effects on expression of large structural variants. These results give us valuable insight into how *M. tb* adapts to new environments. The complexity of the selection pressure applied in this experiment, as well as our focus on characterizing different strains of *M. tb* make these results particularly relevant to the study of natural infections and adaptation of *M. tb* under within-host selection pressures.

## Materials and Methods

### Bacterial strains and growth conditions

Six clinical populations (MT31, MT49, MT55, MT72, MT345, and MT540) were initially isolated from sputum samples and passaged repeatedly as pellicle biofilms as previously described (*17*). Ancestral (prior to biofilm passaging) and evolved (after biofilm passaging) populations were grown both as planktonic cultures and biofilms for RNA sequencing (Figure 1). Planktonic cultures of *M. tuberculosis* were inoculated from freezer stocks into 7H9OTG (Middlebrook 7H9 broth [HiMedia, #M198] containing 0.2% w/v glycerol, 10% v/v OADC supplement [oleic acid, albumin, D-glucose, catalase; Becton Dickinson, # B12351] and 0.05% w/v Tween-80) and incubated at 37^0^C with 5% CO_2_ to an OD600 ∼1 before RNA extraction. For biofilm growth, 250 μL of planktonic culture was inoculated into 25mL Sauton’s medium (for 1L: 0.5g KH_2_PO_4_, 0.5g MgSO_4_, 4g L-Asparagine, 2g Citric acid, 0.05g Ammonium Iron (III) citrate, 60mL glycerol, adjust pH to 7.0 with NaOH) containing 0.1% w/v ZnSO_4_, in a 250 mL flask (Corning, #430281) and incubated at 37^0^C, with 5% CO_2_, without shaking, with a tight-fitting cap. After 3 weeks the lid was loosened to allow gas exchange and the cultures grown for an additional 2 weeks (for a total of 5 weeks of growth) before RNA extraction.

### RNA extraction

Biofilm samples were harvested by first removing and discarding the media below the biofilm. The biofilm was resuspended in fresh Sauton’s medium, pelleted at 5,000 xg for 10 minutes, then resuspended in 3 mL of RNAprotect for bacterial cells (Qiagen, #76506). Planktonic cultures were similarly pelleted and resuspended in RNAprotect. Aliquots (biological replicates) of the RNAprotect suspensions were frozen at -80°C before extraction. Before RNA extraction, RNAprotect suspensions were thawed and pelleted at 5,000 xg for 10 minutes before discarding the supernatant. The pellet was then resuspended in 100 μL of 100 mg/mL lysozyme (Sigma, #L7651) and 150 μL of Proteinase K (Qiagen, #19131), vortexed, and incubated at 37°C for 30 minutes (vortexed every 10 minutes). The remainder of the extraction was performed with a modified version of the Illustra RNAspin mini RNA isolation kit (GE Healthcare, #25-0500-71) protocol: 350 μL of buffer RA1, 3.5 μL of *β*-mercaptoethanol (Sigma, #M6250) and 25-50 mg of acid-washed glass beads (Sigma, #G8772) were added to the cell suspension. Tubes were vortexed at top speed for a total of 3 minutes (6 x 30 sec vortexes with 1 minute ice rest between vortexes) and then centrifuged at 11,000 xg for 1 minute. The supernatant was removed before proceeding with step 3 of section 7.3 of the kit protocol. RNA quantification was performed by Qubit using the RNA HS Assay Kit (Invigrogen, #Q32852).

### Library preparation & sequencing

Total RNA submitted to the University of Wisconsin-Madison Biotechnology Center was assayed for purity and integrity via the NanoDrop One Spectrophotometer and Agilent 2100 Bioanalyzer, respectively. For library preparation, ribosomal RNA was removed from total RNA using the RiboMinus Eukaryote System v2 (ThermoFisher, #A15026) kit following the directions provided by the manufacturer. Using the TruSeq Stranded RNA kit (Illumina, Inc., Carlsbad, CA), the ribo-depleted RNA was fragmented using divalent cations under elevated temperature. Fragmented RNA was copied into first stranded cDNA using SuperScript II Reverse Transcriptase (Invitrogen, Carlsbad, California, USA) and random primers. Second strand cDNA was synthesized using a modified dNTP mix (dTTP replaced with dUTP), DNA Polymerase I, and RNase H. Double-stranded cDNA was cleaned up with AMPure XP Beads (1.8X) (Agencourt, Beckman Coulter). The cDNA products were incubated with Klenow DNA Polymerase to add a single ‘A’ nucleotide to the 3’ end of the blunt DNA fragments. Unique dual indexes (UDI) were ligated to the DNA fragments and cleaned up with two rounds of AMPure XP beads (0.8X). Adapter ligated DNA was amplified by PCR and cleaned up with AMPure XP beads (0.8X). Final libraries were assessed for size and quantity using an Agilent DNA1000 chip and Qubit dsDNA HS Assay Kit (Invitrogen, Carlsbad, California, USA), respectively. Libraries were standardized to 2nM. Paired-end 150bp sequencing was performed, using standard SBS chemistry on an Illumina NovaSeq6000 sequencer. Images were analyzed using the standard Illumina Pipeline, version 1.8.2.

### Custom annotation file

A custom annotation file (available at github.com/myoungblom/mtb_ExpEvo_RNA) was made using the standard annotations for reference strain H37Rv (NCBI accession GCA_000195955.2). To these annotations we added non-coding RNA (ncRNA) compiled from recent publications (*25*–*31*) as well as unpublished data from collaborators (*63*). Finally we included subset of the small open reading frames (sORFs) discovered by C. Smith et al., 2022: only sORFs identified in multiple technical replicates using Ribo-RET, that were antisense to or not overlapping any other genes were included (*64*). The resulting annotation file contains a total of 579 ncRNA and 300 sORFs (Supplementary Data 4).

### RNA sequencing data analysis

Raw RNA-sequencing data was processed using an in-house pipeline (code available at github.com/myoungblom/RNAseq). Briefly, raw data was checked for quality with FastQC v0.11.8 (http://www.bioinformatics.babraham.ac.uk/projects/fastqc) and trimmed using Trimmomatic v0.39 (*65*). Trimmed reads were mapped to the H37Rv reference genome (NCBI accession NC_000962.3) using BWA-MEM v0.7.17 (*66*), and Samtools v1.17 (*67*) view and sort were used to process SAM and BAM files. Assembly quality was determined using Qualimap v2.2.1 BamQC and RNAseq tools (*68*). Finally, gene expression was counted using HTSeq counts v1.99.2 (*18*) using the custom annotation file described above with flags ‘—nonunique none’ and ‘-s reverse’ to exclude reads mapping to more than one feature and to indicate a reverse-stranded library prep, respectively.

### Differential expression analysis

Differential expression analysis was performed in R v4.2.2 (*69*) using DESeq2 v1.30.0 (*19*). First, gene expression counts from HTSeq were loaded using the function ‘DESeqDataSetFromHTSeqCount’, and genes with ‘0’ counts in all samples were removed. A regularized-log transformation was applied to the raw expression counts using the function ‘rlog’ which normalizes counts with respect to library size – these counts were used for visualizations including PCA plots and heatmaps. Significantly differentially expressed genes (DEGs) were identified for each comparison using the function ‘DESeq’, filtering results by *p*-value (with Benjamini-Hochberg correction) of < 0.05. For analyses of all populations, all samples were analyzed with DESeq together using either the growth condition (biofilm or planktonic) or the genotype (evolved or ancestral) as the design factor. For analyses of individual populations, only samples from a given population were included in the DESeq analysis, using the same design criteria as listed above. Scripts for differential expression analyses and figures are available here: github.com/myoungblom/mtb_ExpEvo_RNA.

### Phylogenetic tree

Phylogenetic tree of ancestral populations was inferred using RAxML v8.2.3 as described previously (*17*) and plotted using ggtree (*70*).

### Sliding coverage plots

Sliding window DNA and RNA sequencing coverage was calculated from sorted BAM files created from the mapping pipeline described above, using Samtools bedcov (*67*) with a window size of 20,000 bp and a step size of 5,000 bp (code available at github.com/myoungblom/mtb_ExpEvo). Relative coverage was calculated by dividing each window coverage by the average coverage across the assembly – as calculated by Qualimap BamQC (*68*).

## Data Availability

All RNA sequencing data used for these analyses has been deposited to the NCBI SRA under accession PRJNA927269. Code for processing of raw RNA sequencing data available at github.com/myoungblom/RNAseq. Expression counts and differential expression analysis scripts are available at github.com/myoungblom/mtb_ExpEvo_RNA.

## Acknowledgements

MAY was funded by National Science Foundation Graduate Research Fellowship Program under grant No. DGE-1747503. This work was supported by NIH grant R01-AI113287. The funders had no role in study design, data collection and interpretation, or the decision to submit the work for publication.

The authors thank the University of Wisconsin-Madison Biotechnology Center for sequencing services, and Dr. Sarah Fortune’s lab for their assistance in compiling a list of non-coding RNA.

## Author Contributions

MAY: Data curation, formal analysis, methodology, software, visualization, writing – original draft, writing – review & editing

TMS: Conceptualization, data curation, investigation, writing – review & editing

CSP: Conceptualization, funding acquisition, project administration, supervision, writing – original draft, writing – review & editing

## Supplement

**Table S1:** RNA sequencing sample information. Passage number 1 refers to the ancestral population and evolved populations were sequenced after either 8 or 12 passages. Number of mapped reads and percentage of reads assigned to a feature were calculated using Qualimap v2.2.1(*68*).

| Sample Number | Population | Passage Number | Condition | RNA sequenced (ug) | Mapped reads (millions) | Reads assigned to feature (millions) |
| --- | --- | --- | --- | --- | --- | --- |
| 1 | MT31 | 1 | Planktonic | 1.29 | 73 | 66 |
| 2 | MT31 | 1 | Planktonic | 1.62 | 55 | 50 |
| 3 | MT31 | 12 | Planktonic | 7.04 | 62 | 36 |
| 4 | MT31 | 12 | Planktonic | 1.58 | 53 | 50 |
| 5 | MT31 | 1 | Biofilm | 0.95 | 51 | 32 |
| 6 | MT31 | 1 | Biofilm | 0.98 | 63 | 55 |
| 7 | MT31 | 12 | Biofilm | 2.64 | 79 | 54 |
| 8 | MT31 | 12 | Biofilm | 3.18 | 99 | 39 |
| 10 | MT49 | 1 | Planktonic | 1.52 | 69 | 64 |
| 11 | MT49 | 12 | Planktonic | 3.42 | 76 | 73 |
| 12 | MT49 | 12 | Planktonic | 2.49 | 69 | 56 |
| 13 | MT49 | 1 | Biofilm | 1.54 | 52 | 30 |
| 14 | MT49 | 1 | Biofilm | 3.21 | 117 | 59 |
| 15 | MT49 | 12 | Biofilm | 2.19 | 70 | 34 |
| 16 | MT49 | 12 | Biofilm | 0.95 | 83 | 29 |
| 17 | MT55 | 1 | Planktonic | 1.5 | 69 | 64 |
| 18 | MT55 | 1 | Planktonic | 0.82 | 54 | 50 |
| 19 | MT55 | 8 | Planktonic | 0.44 | 64 | 57 |
| 20 | MT55 | 8 | Planktonic | 3.4 | 75 | 72 |
| 21 | MT55 | 1 | Biofilm | 0.39 | 73 | 70 |
| 22 | MT55 | 1 | Biofilm | 7.33 | 82 | 81 |
| 23 | MT55 | 8 | Biofilm | 1.15 | 75 | 31 |
| 24 | MT55 | 8 | Biofilm | 1.40 | 77 | 32 |
| 25 | MT72 | 1 | Planktonic | 3.15 | 43 | 38 |
| 26 | MT72 | 1 | Planktonic | 1.89 | 62 | 54 |
| 27 | MT72 | 8 | Planktonic | 1.07 | 70 | 58 |
| 28 | MT72 | 8 | Planktonic | 0.72 | 55 | 52 |
| 29 | MT72 | 1 | Biofilm | 1.73 | 52 | 49 |
| 30 | MT72 | 1 | Biofilm | 4.22 | 63 | 54 |
| 31 | MT72 | 8 | Biofilm | 2.24 | 64 | 39 |
| 32 | MT72 | 8 | Biofilm | 1.16 | 54 | 33 |
| 33 | MT345 | 1 | Planktonic | 0.60 | 71 | 58 |
| 34 | MT345 | 1 | Planktonic | 1.14 | 72 | 69 |
| 35 | MT345 | 8 | Planktonic | 1.8 | 73 | 71 |
| 36 | MT345 | 8 | Planktonic | 0.87 | 51 | 49 |
| 37 | MT345 | 1 | Biofilm | 1.05 | 87 | 35 |
| 38 | MT345 | 1 | Biofilm | 1.32 | 87 | 28 |
| 39 | MT345 | 8 | Biofilm | 0.85 | 68 | 28 |
| 40 | MT345 | 8 | Biofilm | 1.2 | 90 | 34 |
| 41 | MT540 | 1 | Planktonic | 0.54 | 69 | 58 |
| 42 | MT540 | 1 | Planktonic | 1.56 | 71 | 68 |
| 43 | MT540 | 8 | Planktonic | 2.73 | 51 | 48 |
| 44 | MT540 | 8 | Planktonic | 2.88 | 72 | 65 |
| 45 | MT540 | 1 | Biofilm | 2.67 | 76 | 74 |
| 46 | MT540 | 1 | Biofilm | 1.91 | 61 | 59 |
| 47 | MT540 | 8 | Biofilm | 1.07 | 63 | 20 |
| 48 | MT540 | 8 | Biofilm | 1.46 | 80 | 33 |

**Figure S1:**
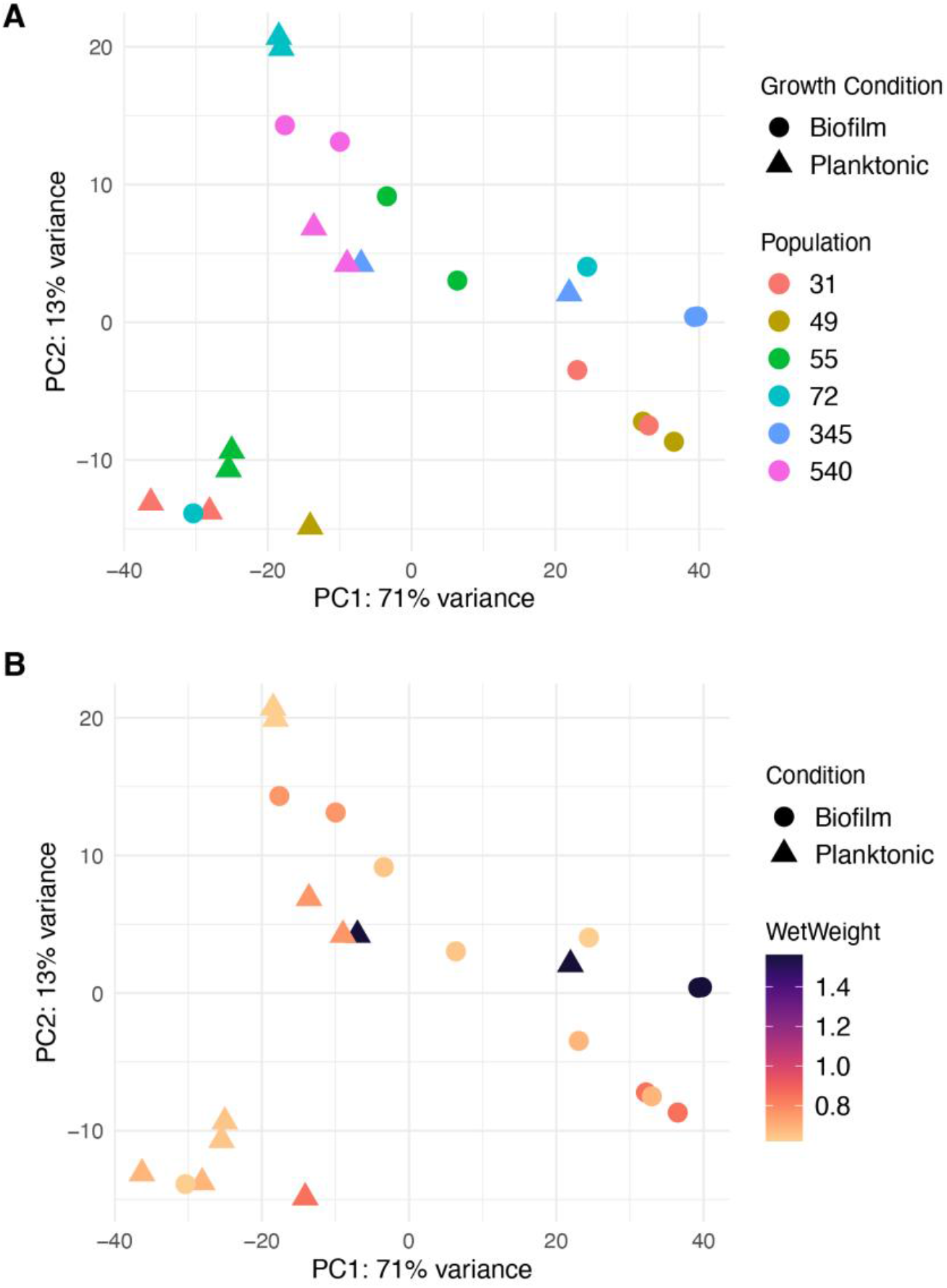
A) Principal component analysis (PCA) of normalized, log-transformed gene expression counts from ancestral populations. Each point represents the total gene expression of a single sample, where the shape indicates the growth condition of the sample, and the color indicates the population. B) PCA (same as in A) colored this time by biofilm wet weight. There is no clear correlation – either among biofilm samples or planktonic samples – between gene expression patterns and ancestral biofilm phenotype (as measured by wet weight).

**Figure S2:**
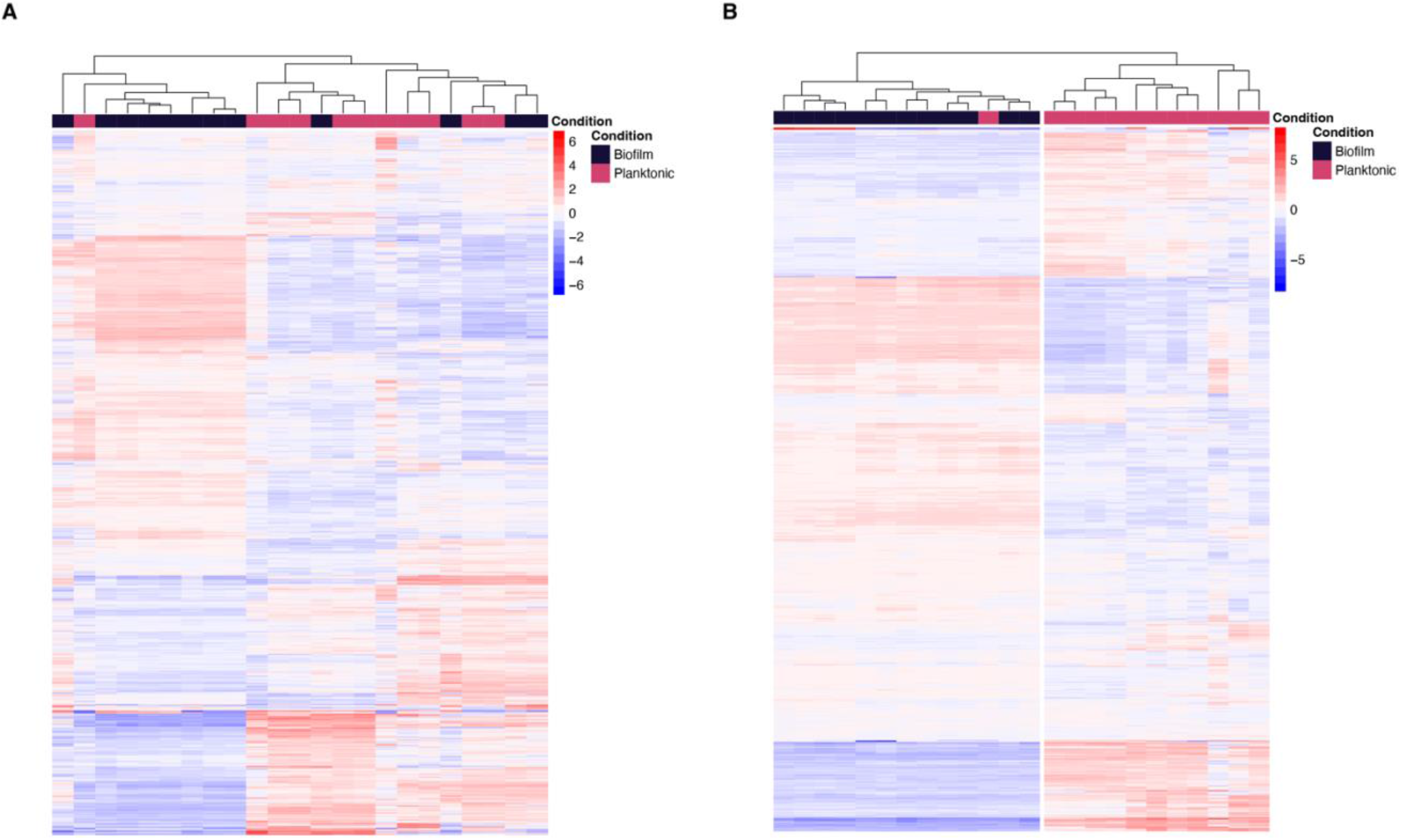
Heatmaps of normalized, log-transformed expression counts all genes in the genome. Each column is a single sample from an ancestral (A) or evolved (B) population, grown either as a biofilm or in a planktonic culture. Each row is a gene. Heatmap colored by expression values for each gene which are normalized to the mean across samples. Samples are clustered by Euclidean distance and plotted as a tree at the top of the heatmap. Evolved populations have much more uniform biofilm transcriptomes.

**Figure S3:**
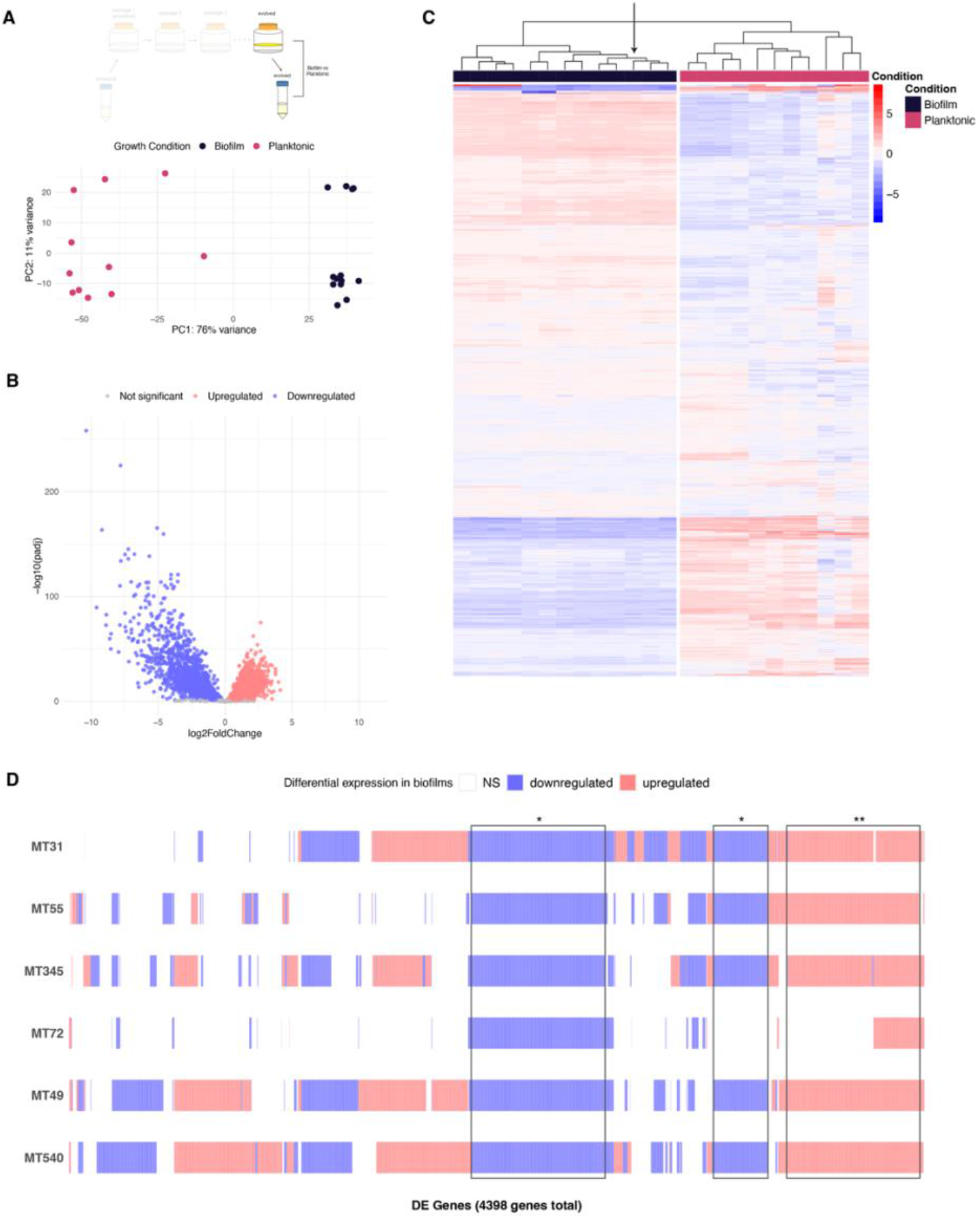
Treating evolved MT31 outlier as a biofilm. A) Top: Experimental diagram highlighting comparator populations: evolved populations grown as pellicle biofilms are compared to the same populations grown in planktonic cultures. Bottom: Principal component analysis (PCA) of normalized, log-transformed gene expression for evolved populations grown as biofilms and planktonic cultures. B) Differential expression of DEGs between evolved populations grown as biofilms and as planktonic cultures. Log transformed adjusted p-values plotted against the log2 fold change for each gene. Genes that did not have significant differential expression are shown in grey. C) Heatmap of normalized, log-transformed expression counts for DEGs shown in panel B. Each column is a single sample from an evolved population, grown either as a biofilm or in a planktonic culture – the sample identified as an outlier in Figure 4 is highlighted by an arrow. Each row is a DEG. Expression values for each gene are normalized to the mean across samples. Samples are clustered by Euclidean distance and plotted as a tree at the top of the heatmap. D) Matrix of individual DEGs shared across evolved populations. A total of 4373 DEGs are plotted according to if that gene is upregulated (red), downregulated (blue) or not significantly differentially expressed (white) in each population. 24% of downregulated genes are shared by at least 5 populations (*), while 16% of upregulated genes are shared by at least 5 populations (**).

**Figure S4:**
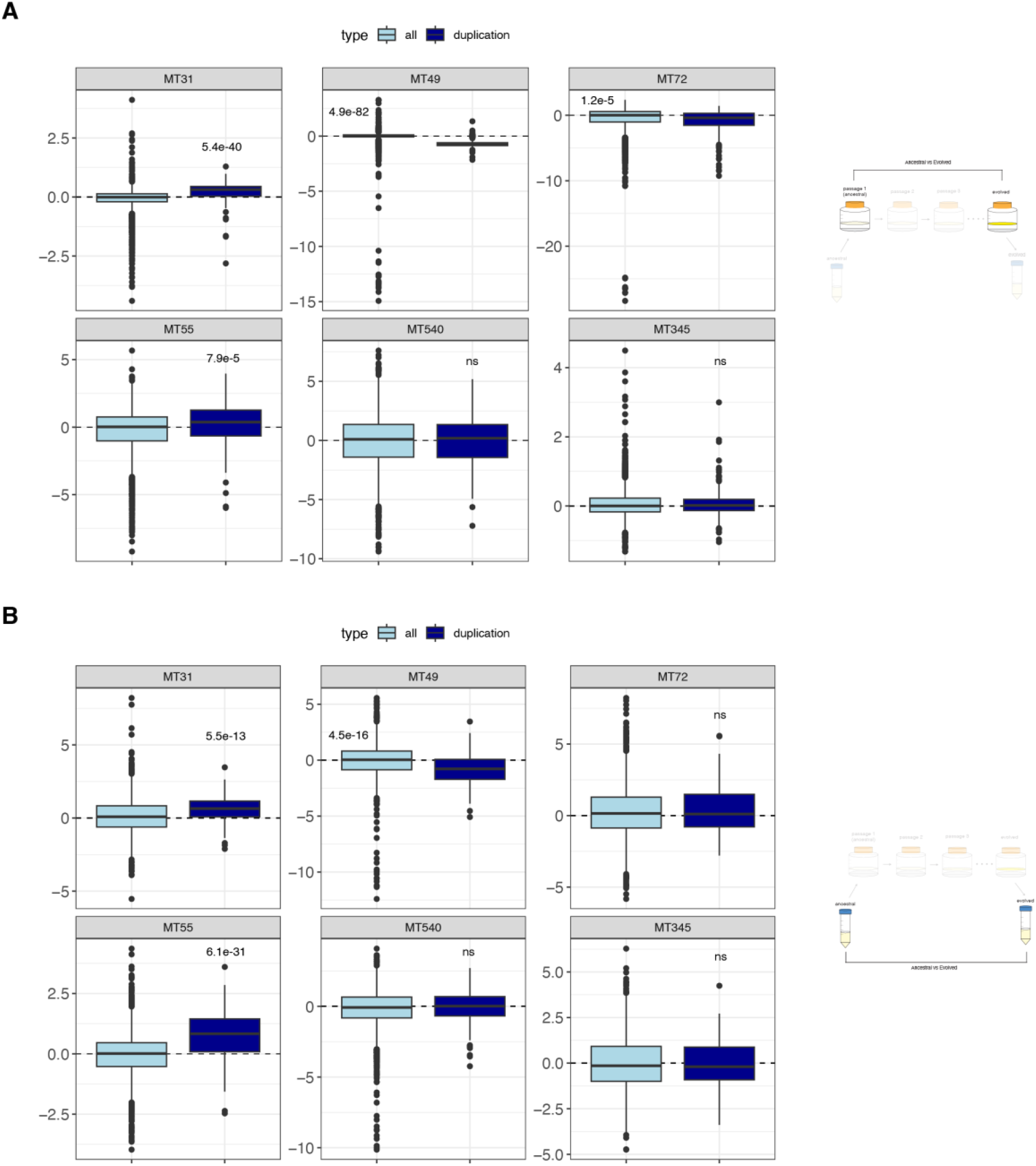
A) Log-2 fold changes (L2FC) for all genes in each biofilm population, separated by their presence in one or both duplications acquired by MT31 and MT55. Mann Whitney U test with Benjamini-Hochberg correction shows significantly higher L2FC values for duplicated genes in MT31 and MT55 (same data shown in Figure 6C), and significantly lower L2FC values for duplicated genes in MT49 and MT72. Comparison is between evolved populations grown as a biofilm and ancestral populations grown as biofilms. B) Same as A, but comparing pellicle evolved populations grown as planktonic cultures, to ancestral populations grown as planktonic cultures. Mann Whitney U test with Benjamini-Hochberg correction shows significantly higher L2FC values for duplicated genes in MT31 and MT55 and significantly lower L2FC values for duplicated genes in MT49.

**Figure S5:**
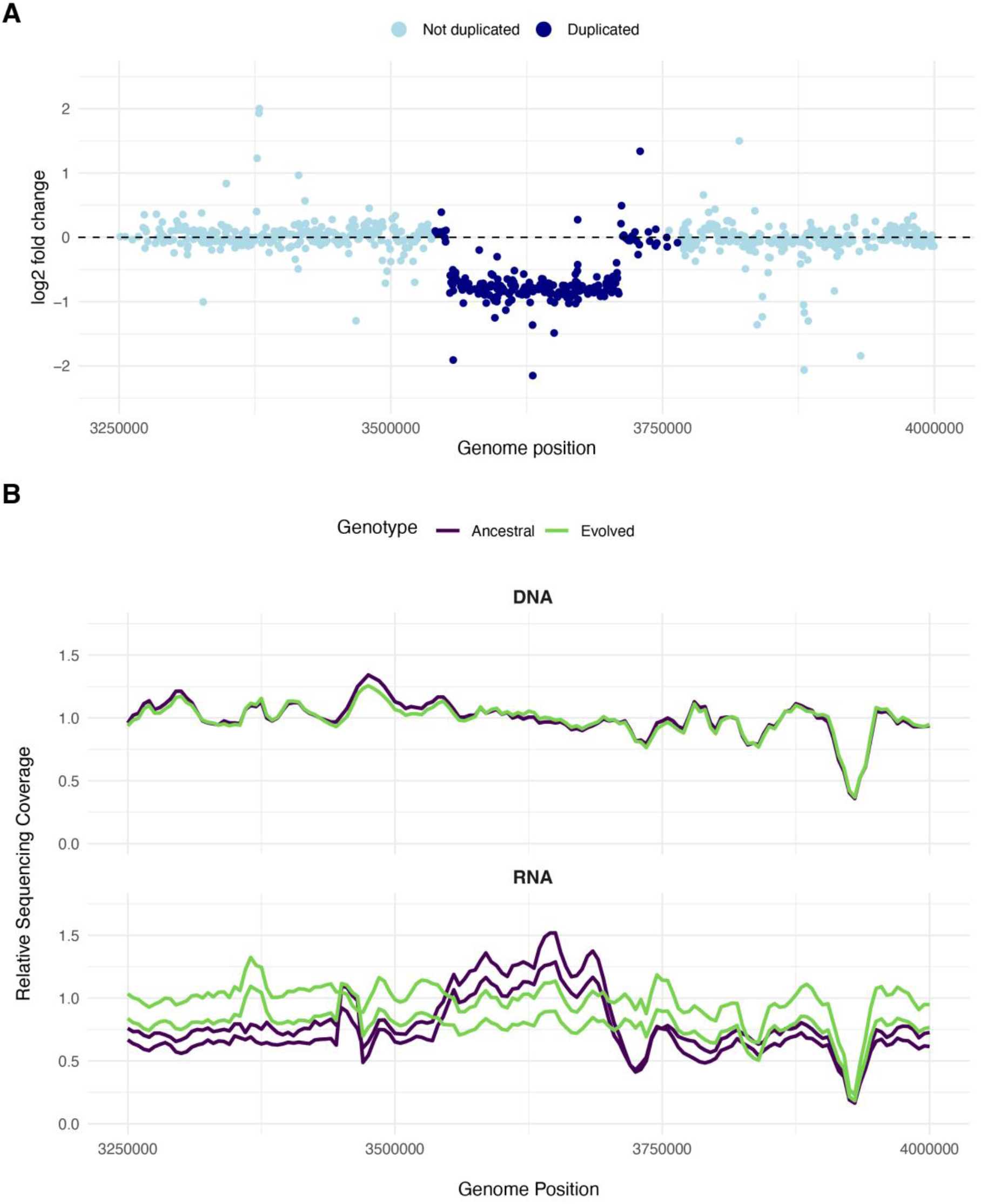
A) log2 fold change (L2FC) values between evolved and ancestral biofilm populations of MT49. Shown is each gene within the region surrounding the duplication which arose in MT31 and MT55. Points are colored according to whether they lie inside of the duplicated region. B) Top: Relative coverage of DNA sequencing for the ancestral (passage 0) and evolved (passage 12) populations of MT49. Bottom: Relative coverage of RNA sequencing for ancestral and evolved populations of MT49. Two lines per genotype indicate two biological replicates.

**Figure S6:**
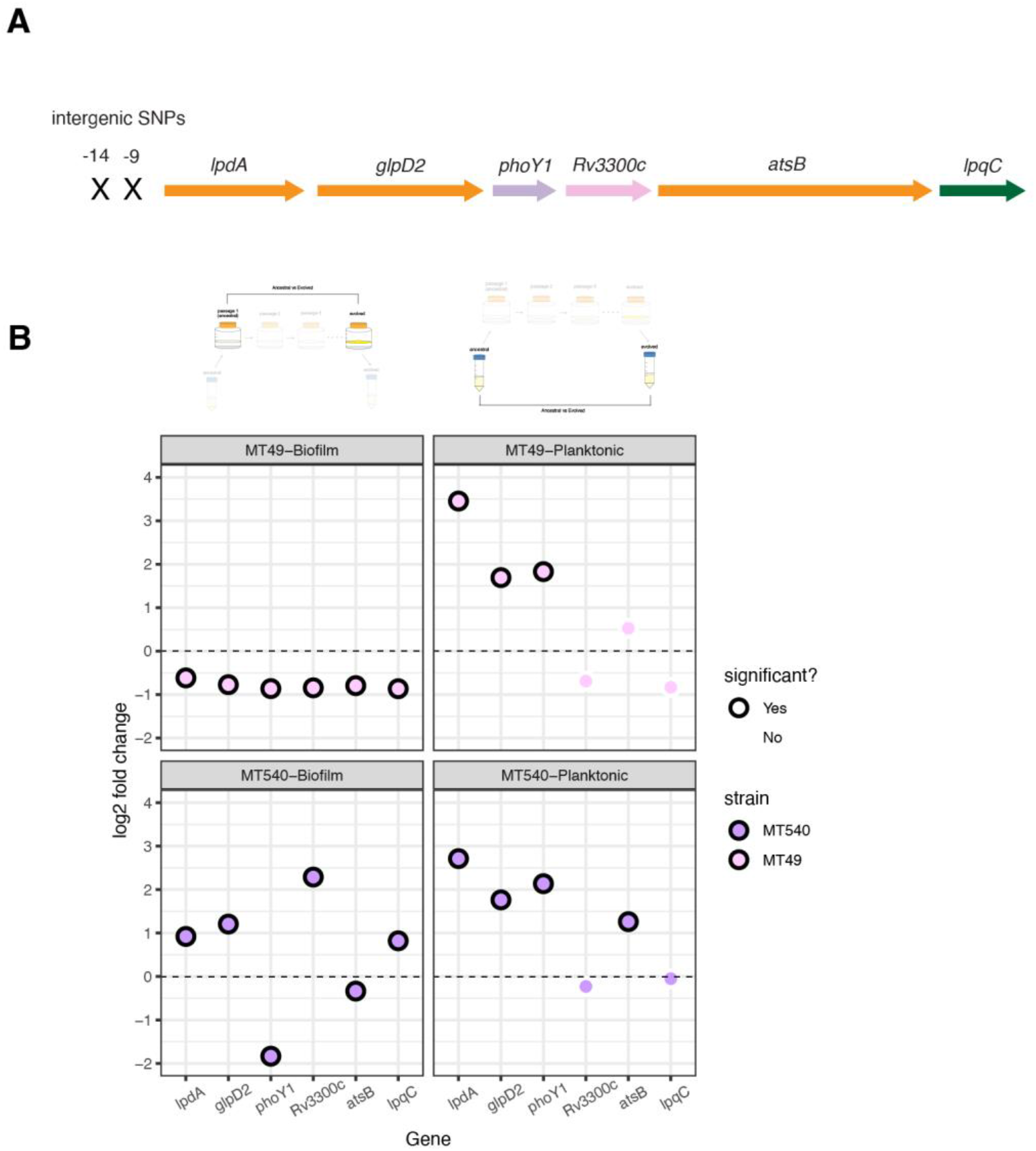
A) Schematic of *lpdA* operon relative to the two intergenic SNPs acquired by MT49 and MT540 over the course of passaging. Genes are colored by functional annotation from Mycobrowser: orange – Intermediary metabolism and respiration, purple – regulatory proteins, pink – conserved hypotheticals, green - cell wall and cell wall processes. B) Log-2 fold change (L2FC) in expression comparing evolved to ancestral populations, for genes downstream of a convergent intergenic mutation in two of our populations (MT49 and MT540). Points circled in black indicate significant L2FCs in expression between evolved and ancestral populations. ‘Biofilm’ refers to evolved populations grown as a biofilm, compared to ancestral populations grown as a biofilm, and the same is true for ‘planktonic’ as shown in diagrams at the top of the panel.

**Figure S7:**
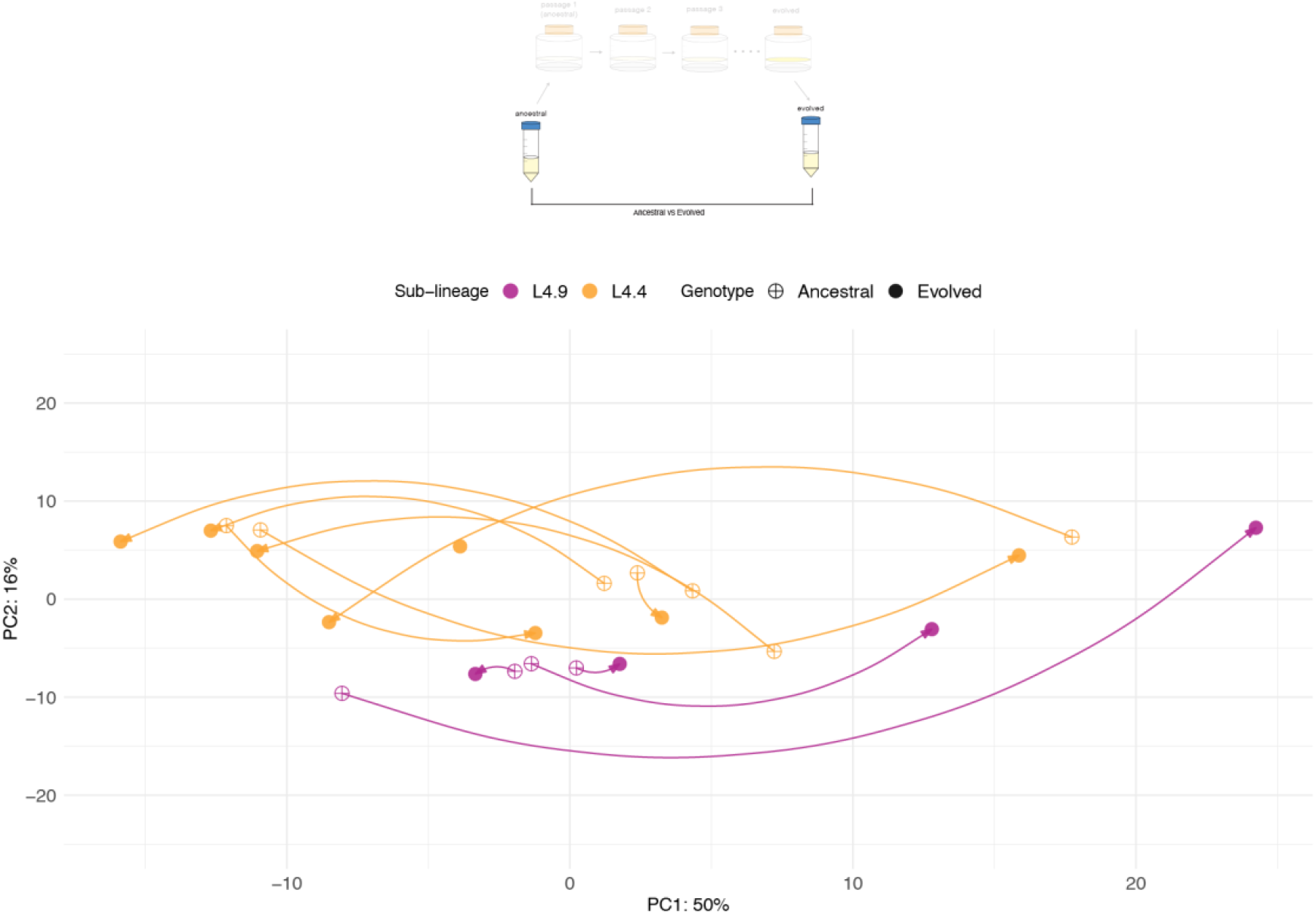
Principal component analysis (PCA) of normalized, log-transformed expression counts of ncRNAs and sORFs from ancestral and evolved populations grown in planktonic culture. Arrows are drawn between corresponding ancestral and evolved populations, indicating the trajectory of evolution across passaging. Points are colored by sub-lineage of the ancestral population.

## Supplementary Data

**SupplementaryData_1.xlsx**: DEGs comparing ancestral populations grown as biofilms, to the same populations grown under planktonic conditions. DEGs provided for multiple analyses: all populations and each population individually.

**SupplementaryData_2.xlsx**: DEGs comparing pellicle passaged (evolved) populations grown as biofilms, to the same populations grown under planktonic conditions. DEGs provided for multiple analyses: all populations and each population individually.

**SupplementaryData_3.xlsx**: DEGs comparing pellicle passaged (evolved) populations grown as biofilms, to ancestral populations grown as biofilms. DEGs provided for multiple analyses: all populations, each population individually and for each of the two sub-lineages in our sample (L4.4 and L4.9).

**SupplementaryData_4.xlsx**: Non-coding RNAs (ncRNAs) and small open reading frames (sORFs) included to make the custom annotation file.

